# Expanding the molecular language of protein liquid-liquid phase separation

**DOI:** 10.1101/2023.03.02.530853

**Authors:** Shiv Rekhi, Cristobal Garcia Garcia, Mayur Barai, Azamat Rizuan, Benjamin S. Schuster, Kristi L. Kiick, Jeetain Mittal

## Abstract

Understanding the relationship between an amino acid sequence and its phase separation has important implications for analyzing cellular function, treating disease, and designing novel biomaterials. Several sequence features have been identified as drivers for protein liquid-liquid phase separation (LLPS), leading to the development of a “molecular grammar” for LLPS. In this work, we further probed how sequence modulates phase separation and the material properties of the resulting condensates. Specifically, we used a model intrinsically disordered polypeptide composed of an 8-residue repeat unit and performed systematic sequence manipulations targeting sequence features previously overlooked in the literature. We generated sequences with no charged residues, high net charge, no glycine residues, or devoid of aromatic or arginine residues. We report that all but one of the twelve variants we designed undergo LLPS, albeit to different extents, despite significant differences in composition. These results support the hypothesis that multiple interactions between diverse residue pairs work in tandem to drive phase separation. Molecular simulations paint a picture of underlying molecular details involving various atomic interactions mediated by not just a handful of residue types, but by most residues. We characterized the changes to inter-residue contacts in all the sequence variants, thereby developing a more complete understanding of the contributions of sequence features such as net charge, hydrophobicity, and aromaticity to phase separation. Further, we find that all condensates formed behave like viscous fluids, despite large differences in their viscosities. The results presented in this study significantly advance the current sequence-phase behavior and sequence-material properties relationships to help interpret, model, and design protein assembly.

## Introduction

Biomolecular condensates are responsible for diverse cellular functions in the cytoplasm and the nucleus^1–4^. They often form through liquid-liquid phase separation (LLPS), thus retaining liquid-like properties^5, 6^. Understanding the role of different amino acids and the associated molecular interactions that drive the thermodynamics and dynamics of phase separation is valuable for gaining a fundamental understanding of these biologically important assemblies^7–10^. Furthermore, such an understanding can help unlock the potential of condensates for a variety of applications ranging from materials^11–14^ to synthetic biology^15–17^.

Recent studies on natural and artificial sequences have painted a picture of the drivers of protein LLPS^11, 18–26^. Many proteins that undergo LLPS are found to be intrinsically disordered or contain intrinsically disordered regions (IDRs), often of low complexity, i.e., enriched in polar, aromatic, and proline and glycine residues^7, 27, 28^. It has become clear from recent studies that higher content and segregation of aromatic residues, often Tyr, can underlie protein phase separation^21, 23, 29, 30^. Additionally, Arg, but not Lys, has been proposed to drive phase separation through interaction with aromatic residues^18, 19, 21, 31^. It is hypothesized that the overall interactions between the Arg and aromatic sidechains – including hydrogen bonding, sp^2^/π, and the electrostatic interaction between the guanidinium group and the aromatic ring – contribute to the strength of this interaction^12, 20, 21, 24^. Many studies have also highlighted the role of charge-charge attraction between oppositely charged residues (Arg/Lys with Asp/Glu) in modulating the phase behavior of different proteins^19, 24, 32^.

Building upon considerable work that has established important molecular features of LLPS^11, 19–22, 24–26, 29^, here we consider what additional interactions play key contributing roles in LLPS and how these interactions might be leveraged to fine-tune LLPS and the physical properties of the condensates. To tackle this, we designed and tested sequences that deviate from the existing heuristics and predictors of phase separation. In conducting these studies, we also addressed another key question: To what extent can protein LLPS be modeled as a process driven by a handful of residues, or is it necessary to account for interactions between many diverse residues?

We began our studies with the sequence (GRGDSPYS)_25_, an artificial intrinsically disordered protein (A-IDP) with upper critical solution temperature (UCST) phase behavior, which was recently studied by Dzuricky et al.^12^ and Dai et al.^33^. We then carefully designed new unique variants of this polypeptide to dissect the role of different amino acids on their own, and in conjunction with other residues, on *in vitro* phase separation. We examined sequences that are uncharged, have a large net charge, lack aromatic or arginine residues, or are devoid of glycines. Such sequences lack sequence features that are believed to be critical based on the current understanding of LLPS^20, 34–36^. Surprisingly, we identified sequences that undergo phase separation in each category to form protein droplets with liquid-like properties. We obtained mechanistic insights into the atomic interactions of these amino acids using state-of-the-art all-atom molecular dynamics simulations. Furthermore, using passive microrheology experiments, we quantified changes in the viscosity of the condensates as a result of specific amino acid substitutions. Collectively, this work expands our understanding of the molecular language of protein LLPS significantly beyond what is known currently.

## 2. Results and Discussion

### 2.1 WT A-IDP undergoes LLPS driven by multivalent interactions involving multiple residues

The first A-IDP we examined here consists of 25 repeats of the 8-residue peptide^12^ GRGDSPYS, denoted in our study as wild-type (WT). The repeat unit contains several interactions believed to play a key role in modulating phase separation, including hydrogen-bonding^24, 37^, electrostatic attraction and repulsion^18, 19, 24^, aromatic-cationic interactions^12, 18, 19, 21, 26^, sp^2^/*π* interactions^20, 21, 26^, and van der Waals interactions^10, 37^ (**Fig. 1a**). Additionally, the A-IDP is similar in length and shows high compositional similarity to proteins from the FET family as well as other naturally occurring prion-like low complexity domains (PLCDs)^27^ (**Fig. 1a**). This similarity of A-IDP composition supports its use to examine the completeness of the existing molecular grammar originally proposed for FUS-like proteins^21^, later extended to PLCDs^26^.

**Fig. 1.**
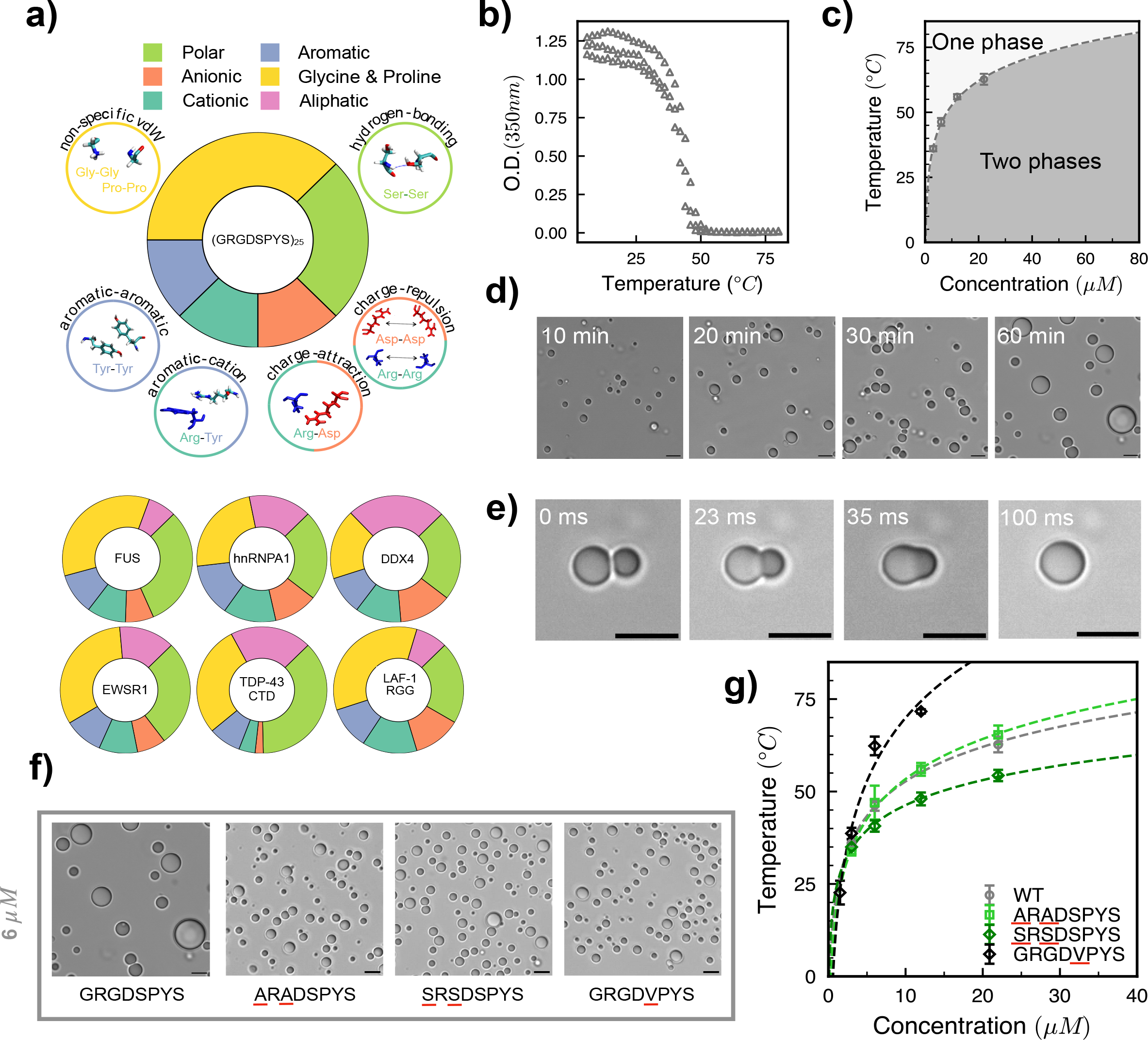
A diverse range of interactions between residue pairs contributes to phase separation of WT, (GRGDSPYS)_25_. (a) Pie charts comparing the composition of the WT polypeptide to naturally occurring sequences. The composition is segregated into Polar; Aromatic; Anionic; Glycine and Proline; Cationic; and Aliphatic residues. Surrounding the WT pie chart are cartoons highlighting the different residue pair interactions present in WT. (b) Example turbidimetry experiments on WT to estimate transition temperatures; shown here are triplicate turbidity assays at WT concentration 6 µM. (c) Partial phase diagram of WT obtained through turbidimetry at different WT concentrations in PBS. The dashed line is obtained through a logarithmic fit to the transition temperatures at each polypeptide concentration. (d) Microscopy images of WT at 6 µM concentration at time points 10, 20, 30, and 60 minutes after inducing phase separation (Scale bar, 5 µm). (e) Images of WT droplets undergoing coalescence over a 100 ms time window (Scale bar, 5 µm). (f) Microscopy images of WT, ARADSPYS, SRSDSPYS, and GRGDVPYS variants at 6 µM concentration (Scale bar, 5 µm). (g) Partial phase diagram of WT, ARADSPYS, SRSDSPYS, and GRGDVPYS variants in PBS obtained through turbidimetry at different concentrations. Dashed lines are obtained through a logarithmic fit to the measured transition temperatures at each concentration from turbidity experiments.

#### 2.1.1 Investigating diverse interactions resulting in A-IDP phase separation

The A-IDP repeat unit, due to its similarity to resilin and other Resilin-Like Polypeptides (RLPs), and based on previous work, is expected to show UCST behavior^12, 38^. Our results from turbidimetry experiments at 6 *μ*M protein concentration show a clear transition at a temperature of 45 °C (**Fig. 1b**) in phosphate-buffered saline (PBS, pH 7.4). Turbidity measurements at several concentrations (**Fig. S1**) are used to generate a partial phase diagram, consisting of the low concentration side (left arm), which is also referred to as the saturation concentration (C_sat_) (**Fig. 1c**). We then investigated whether the turbidity is due to formation of spherical droplets, a characteristic of LLPS – and not due to aggregation – by taking microscopy images at a 6 *μ*M concentration. We observed that WT forms spherical droplets that grow with time when allowed to phase separate at room temperature (**Fig. 1d**). To check whether these spherical droplets are viscous liquids or have liquid-like nature, we monitored droplet fusion events, which confirmed these droplets are contacting, fusing, and rounding into larger spherical droplets (**Fig. 1e**). The aspect ratio vs. time plot for a typical fusion event shows that the relaxation time of these fusing droplets is less than 100 ms, which is comparable to that of similarly sized droplets comprised of the LAF-1 RGG domain and some of its mutants^24^ (**Fig. S2**).

Because the WT lacked aliphatic hydrophobic residues, which are present in other PLCDs (**Fig. 1a**), we produced recombinant Gly-to-Ala (<u>A</u>R<u>A</u>DSPYS)_25_ and Ser-to-Val (GRGD<u>V</u>PYS)_25_ variants. (All sequences described in this study contain an octapeptide repeated 25 times, so for brevity we often drop the subscript 25 hereafter.) Neither variant shows much change in the saturation concentration at room temperature, as compared to WT. However, the Ser-to-Val variant results in a dramatic increase in the transition temperature (T_t_), from 46.3 ± 1.5 °C to 62.3 ± 2.5 °C, at 6 *μ*M concentration. This indicates that GRGD<u>V</u>PYS condensates are more thermally stable, eventually showing a lower saturation concentration than WT at temperatures above ∼40 °C (**Fig. 1g**).

We further find that Gly-to-Ser mutations (<u>S</u>R<u>S</u>DSPYS) lead to comparable LLPS behavior as that of Gly-to-Ala (and WT) at lower concentrations, although reduced transition temperatures are observed at higher concentrations for this variant (**Fig. S3**), highlighted by a downward shift in the phase diagram (**Fig. 1g**). These results suggest that Ala and Gly residues impart similar LLPS behavior in A-IDPs, but the polar Ser residue contribution can be similar or different to Ala/Gly, depending on the solution concentration or temperature. Importantly, microscopy experiments show the liquid-like nature of the droplets formed by all these variants, indicating that Gly residues may not be required for the formation of liquid-like droplets (**Fig. 1f**).

#### 2.1.2 Atomistic simulations highlight diverse interactions within the condensate involving polar residues

We conducted a microsecond-long all-atom explicit solvent molecular dynamics simulation of a WT protein condensate (**Fig. 2a**) using a previously reported strategy^39^ (see Methods) to gain insights into the residue-level molecular interactions stabilizing the protein-rich phase. We find that the protein condensate remains stable over the 1 *μ*s time with rapid equilibration of solvent density and a relatively slow change in protein concentration profile over this time (**Fig. S4**). The average protein density after removing the initial 250 ns equilibration is 428 mg/ml (**Fig. 2b**), which is similar to other protein condensates such as FUS LC (∼447 mg/ml)^22^ and DDX4 (∼400 mg/ml)^19^. Due to the presence of a zwitterionic pair (Arg and Asp residues) in the polypeptide repeat unit, the concentrations of Na and Cl ions are similar to each other both inside and outside the condensate (**Fig. 2b**). Interestingly, the concentration of Na and Cl ions is lower inside the protein condensate by a factor of two, reflecting the free energy penalty associated with the desolvation of ions^39^ .

**Fig. 2.**
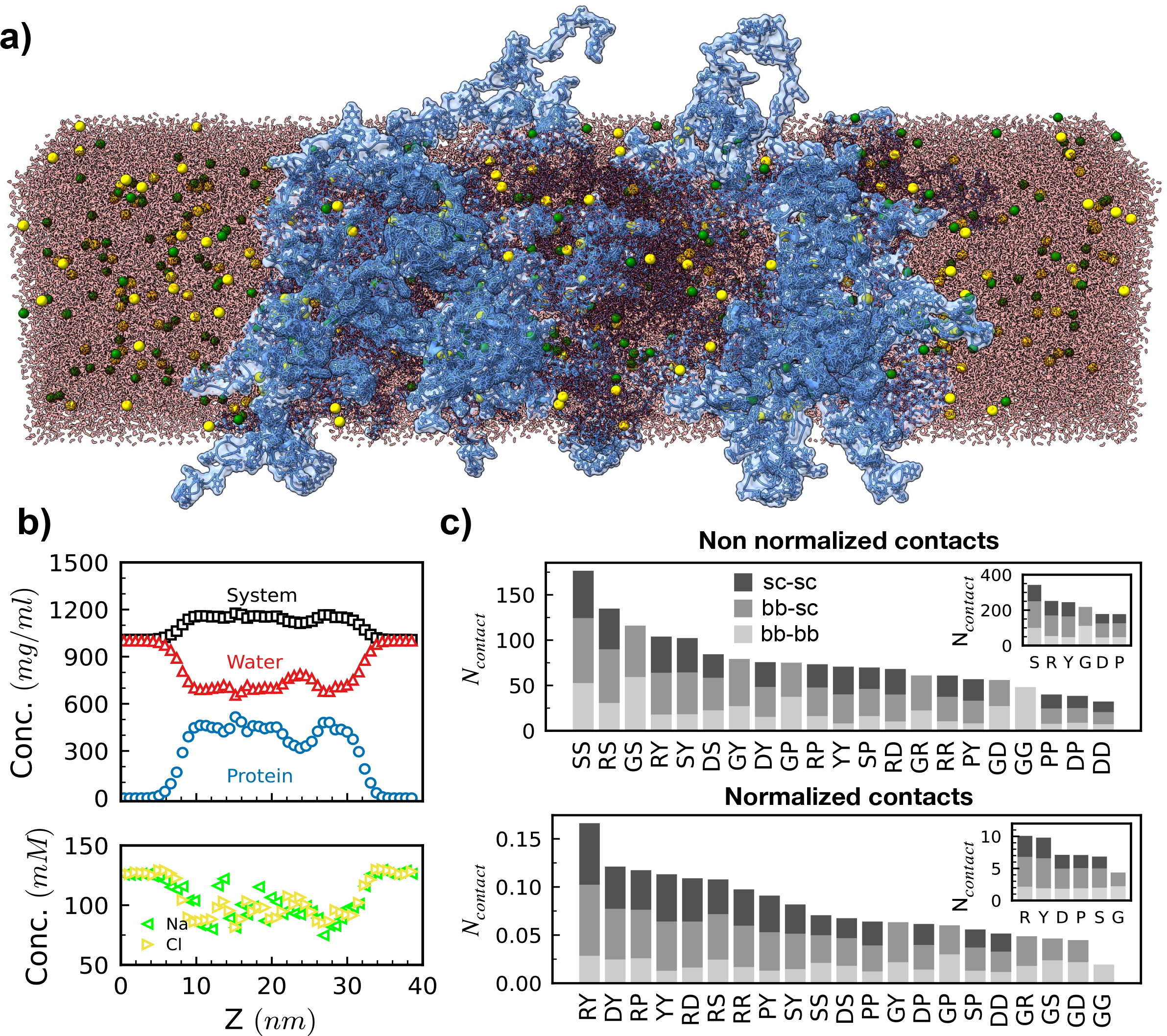
Atomistic simulations of the WT sequence highlight the diverse interactions encoded within the sequence. (a) Representative snapshot of the atomistic simulation of the condensed phase of WT. Proteins are shown as a semitransparent surface with bonds and atoms shown explicitly in blue; water in red; and Na and Cl ions in green and yellow, respectively. (b) Estimated densities of system (protein, water, and ions combined), and of water, protein, sodium ions, and chlorine ions from atomistic simulations of the WT condensed phase, consisting of 30 protein chains in the slab geometry. (c) Average residue pair contacts estimated from atomistic simulations of the WT condensed phase, separated into backbone-backbone (bb-bb), backbone-sidechain (bb-sc) and sidechain-sidechain (sc-sc). The inset shows the average contacts formed by each residue during the simulation. The bottom panel shows the average pair-wise contacts formed by the residues when normalized by their abundance within the WT sequence, and the inset shows the individual residue contacts with the same normalization applied.

Next, we used the simulation data to dissect the prevalence of specific amino acid pairings (such as Arg:Tyr, Tyr:Tyr, and Arg:Asp) inside the protein condensate, by calculating the average number of contacts formed by different residue pairs. Consistent with previous computational work on the FUS low-complexity (LC) domain and LAF-1 RGG domain^39^ and NMR experiments of FUS LC^22, 25^, we find that effectively all residue pairs form contacts through backbone-backbone (bb-bb), sidechain-sidechain (sc-sc), and backbone-sidechain (bb-sc) atomic interactions (**Fig. 2c top**). Perhaps somewhat surprisingly in the context of the current understanding of LLPS molecular grammar, we find that contacts between Ser residues (S:S) are observed most frequently, followed by contacts formed by Ser with Arg (S:R) and Gly (S:G). The contacts between aromatic Tyr with Arg (R:Y) and with itself (Y:Y) are also highly prevalent. In fact, if we sum the number of contacts formed by a specific residue (**Fig. 2c top, inset**), Ser forms the highest number of contacts, followed by Arg, Tyr, Gly, Asp, and Pro. As Ser and Gly appear twice as many times as any other residue in the WT sequence (GRGDSPYS), we believe that their higher contact tendency simply reflects their higher composition, because this calculation doesn’t account for the quantitative differences in the interaction strengths between smaller (Ser/Gly) versus bulkier (Arg/Tyr) residues.

In fact, normalizing the number of contacts by amino acid frequency highlights that contacts formed by Tyr and Arg residues with each other and with other residues are the most populated (**Fig. 2c bottom**), which is in line with the current expectation that these residues act as primary drivers of LLPS^18, 19, 21, 26^. To rule out the possibility that the commonly used restrictive distance-based contact definition is overcounting contact pairs involving non-Arg/Tyr residues^32^, we estimate the energetic contribution of different contact pairs (see Methods and Ref. for details). We find that residue pair contacts apart from those between oppositely charged residues Arg and Asp (R:D), i.e., involving Gly, Ser, and Tyr, are also energetically favorable (**Fig. S5**). Together, the results of atomistic simulations strongly support the relevance and contributions of interactions mediated by uncharged, non-aromatic residues in stabilizing protein condensates.

### 2.2 Removal of zwitterionic pair reduces phase separation propensity

To further investigate the contribution of non-electrostatic interactions to LLPS, we tested mutations targeting the zwitterionic pair in the WT repeat unit. We began by replacing Arg and Asp residues with polar residues, Gln and Asn, respectively. Notably, the new variant, (G<u>Q</u>G<u>N</u>SPYS)_25_, also undergoes LLPS at a concentration of 6 *μ*M, albeit with a lower transition temperature of 25.3 ± 0.6 °C (versus 46.3 ± 1.5 °C for WT) at this concentration (**Fig. 3a**, **Fig. S6**). The reduced LLPS propensity of this variant due to the removal of favorable attractions of Arg with Asp (electrostatic attraction) and Tyr (e.g., cation-π) is also highlighted by a downward shift in the phase diagram and increase of *C_sat_* at a fixed temperature (**Fig. 3a**). As with WT, the formation of dense liquid-like droplets and growth in droplet size with time is observed via microscopy of (G<u>Q</u>G<u>N</u>SPYS)_25_ (**Fig. 3b**, **Fig. S7**). Despite losing favorable interactions upon mutation of Arg and Asp, the polypeptide variant without the zwitterionic pair can still undergo LLPS at relatively low concentrations (**Fig. 3c**), with *C_sat_* ∼ 4 μM at 20 *°*C, as compared to FUS LC *C_sat_* ∼125 µM at physiological salt conditions and hnRNPA1 *C_sat_* ∼100 *μ*M at 20 °C.^21, 26^ This result supports the hypothesis that diverse interactions drive LLPS in the WT polypeptide, as identified by our atomistic simulations.

**Fig. 3.**
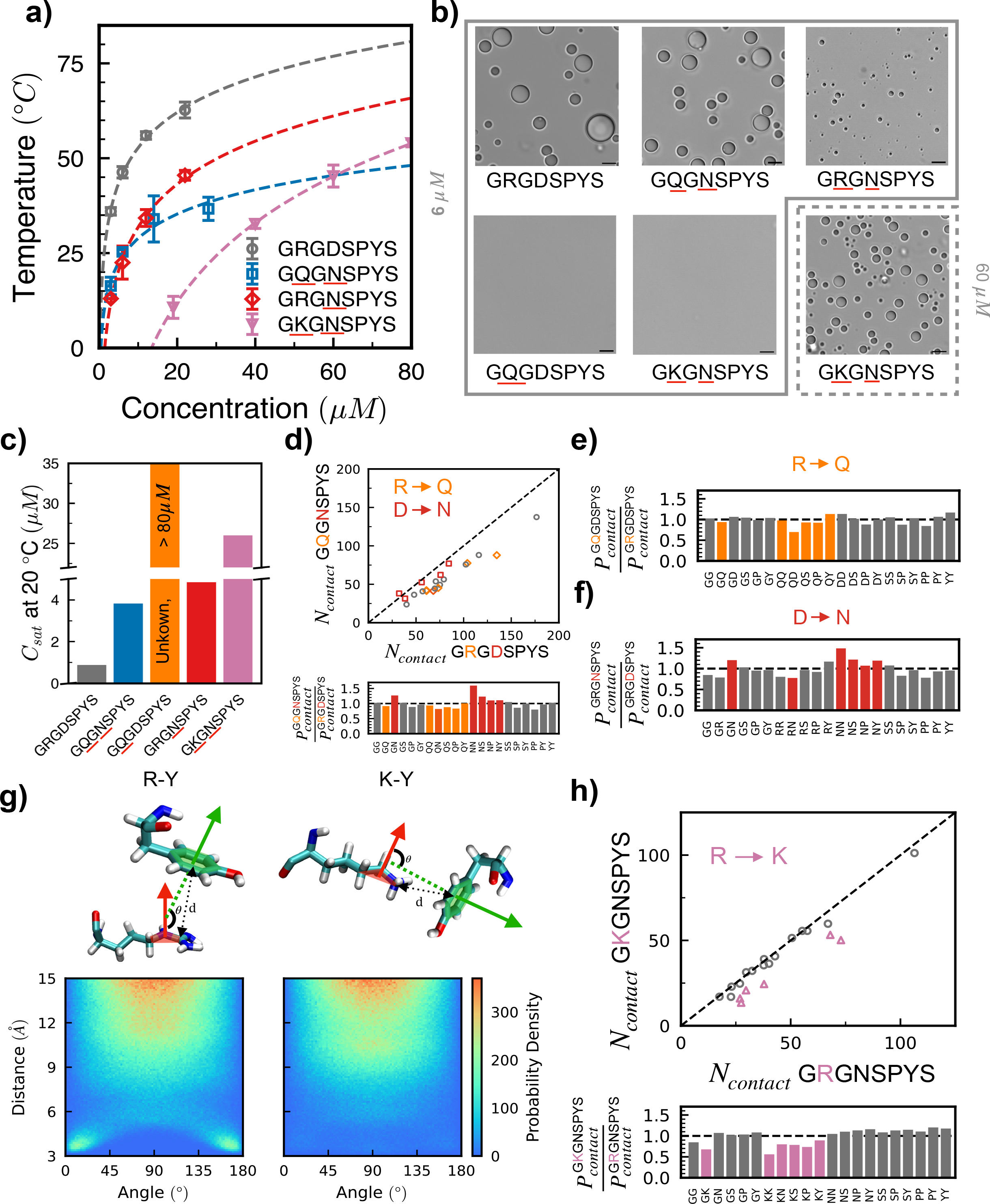
Presence of arginine promotes but is not required for phase separation. (a) Partial phase diagrams for WT (GRGDSPYS), GQGNSPYS, GRGNSPYS, and GKGNSPYS in PBS. GQGDSPYS shows no measurable transition even up to a concentration of 80 µM and is thus omitted from the plot. Dashed lines are obtained through a logarithmic fit to the measured transition temperatures from turbidity experiments at different polypeptide concentrations. (b) Microscopy images for the WT and the different variants at a concentration of 6 µM, shown in the box bounded by solid lines, and additionally at 60 µM for the GKGNSPYS variant in the box bounded by dashed lines. (Scale bar, 5 µm) (c) Saturation concentrations (C_sat_) measured at 20 °C for the different variants. GQGDSPYS shows no measurable transition even up to a concentration of 80 µM. (d) Average residue pair-wise contacts estimated through atomistic simulations for GQGNSPYS. Residue pair-wise contacts are plotted with respect to WT. The diagonal indicates equal number of contacts in WT and the variant. Residue pairs not involving the mutated residues are shown as gray circles, while residue pairs involving the mutated residues are shown in accordance with the color code for the mutations in the plot (R to Q in orange, D to N in red). The lower plot shows the ratio of the probability of contact formation between residue pairs in the variant and the WT. The bars follow the same color code for mutations. (e,f) Ratio of probability of contact formation between residue pairs in the GQGDSPYS (e) and GRGNSPYS (f) variants and the WT. Contacts not involving the mutated residue are shown as gray bars, while contacts involving the mutated residue are shown in orange (R to Q) or red (D to N). (g) Angle vs. Distance plots highlighting the frequency of occurrence of different configurations adopted between Tyr and Arg (left) and Lys (right) in atomistic simulations of the condensed phases in the slab geometry of the GRGNPYS and GKGNSPYS variants. The snapshot above the respective plots shows the definition of the measured angle denoted by θ, and distance, denoted by *d*. (h) Correlation plot similar to (d), comparing residue pair contacts between GRGNSPYS and GKGNSPYS variants. Contacts involving mutated R to K residues are shown in magenta

We simulated this new variant by introducing Arg-to-Gln and Asp-to-Asn mutations in the final configuration of the WT system (see Methods for details) and find that the protein condensate remains stable throughout the 1 *μ*s simulation time (**Fig. S8**). Similar to WT, we observe interactions between most amino acid pairs inside the condensed phase (**Fig. S9**). When comparing the number of contacts between the WT and the G<u>Q</u>G<u>N</u>SPYS variant (**Fig. 3d**), an overall reduction in contacts is observed upon removing the zwitterionic pair (with a notable exception for contacts involving the Asn residue), which is consistent with the variant’s reduced phase separation in experiments. To reveal changes in the pairwise contact formation in a manner that accounts for differences between sequences in their total number of contacts, we normalize *P_contact_* for the variant with *P_contact_* of WT, where *P_contact_* = *N_contact_* / *N_total_* and *N_total_* is the total number of contacts formed by all residue pairs. We observe lower contact probability for pairs involving Arg-to-Gln mutations, while many other pairs (e.g., G:G, G:Y, S:S, S:Y, P:Y, and Y:Y) show no appreciable change, further highlighting the need to consider uncharged, non-aromatic residues and their contribution to LLPS (**Fig. 3d**).

### 2.3 Polycationic and polyanionic variants exhibit contrasting phase behavior

Next, we sought to resolve outstanding questions regarding interactions involving Arg and its contributions to LLPS. We replaced only Asp with Asn (yielding repeat unit GRG<u>N</u>SPYS) to interrogate the role of Arg in LLPS of a polypeptide containing aromatic Tyr but devoid of anionic residues (i.e., in the absence of electrostatic attraction)^11, 36, 40, 41^. Remarkably, at 6 µM, we observe formation of droplets with liquid-like material properties for this highly charged polycationic variant (**Fig. S7**, **Fig. 3a, b**), though the transition temperature is significantly reduced with respect to WT and is slightly reduced from G<u>Q</u>G<u>N</u>SPYS (**Fig. S6**). It is worth noting that the *C_sat_* of this polycationic variant is slightly higher than G<u>Q</u>G<u>N</u>SPYS at lower temperatures, but its *C_sat_* is lower at higher temperatures (**Fig. 3a and Fig. 3c**), showing the need to consider environmental conditions, such as temperature, salt, etc., as part of the complete molecular language of LLPS.

We also tested a polyanionic variant in which we replaced only Arg with Gln (G<u>Q</u>GDSPYS). The turbidity measurements suggest no phase separation even up to high concentrations of 80 *μ*M (**Fig. S6**), which is further confirmed by the lack of droplet formation from the microscopy experiments (**Fig. 3b and Fig. S7**). This behavior contrasts with that of the polycationic variant containing Arg, but it is consistent with expectation for reduced phase separation based on the high net charge of the polyanion and in the absence of crowding agents or elevated salt concentration^42^.

Atomistic simulations of the condensates formed by the polycationic and polyanionic sequences show a significant expansion of the protein-rich phase within the simulation box (**Fig. S8**), and the protein-rich phase incorporates a much higher number of oppositely charged ions to neutralize the excess charge. The number of contacts formed between all residue pairs is lower in these variants than in WT (**Fig. S9, Fig. S10**), but the normalized data again highlight the prevalence of contacts between all other residues pairs (**Fig. 3e,f**, **Fig. S9**).

### 2.4 Polycationic sequence with Arg-to-Lys substitutions still undergoes phase separation

Due to the seminal work of Wang et al.^21^ on FUS-family proteins, it is widely believed that attractive contacts between Arg and Tyr are indispensable to the LLPS of proteins more generally that contain these residues. However, a simple empirical theory to predict *C_sat_* based on the number of Arg and Tyr residues does not capture the observed behavior for many proteins^24^. We therefore revisited the role of Arg and Tyr in the context of the A-IDP sequences. Cation-π interactions^43^, through quantum chemical calculations, have been shown to be much weaker in aqueous solution for Lys than Arg^44–46^, so we thus carried out Arg-to-Lys substitutions in the polycationic sequence to test this directly. The new variant (G<u>K</u>G<u>N</u>SPYS)_25_ does not display LLPS at 6 *μ*M (**Fig. S6**, **Fig. 3b**), although with an increase in concentration of just ∼3-fold (to 20 *μ*M), this Lys-rich polycationic solution becomes turbid below a transition temperature of 10.7 ± 2.9°C (**Fig. S6**). Microscopy observation confirms that (G<u>K</u>G<u>N</u>SPYS)_25_ assembles into liquid-like droplets at room temperature for 60 *μ*M protein concentration (**Fig. 3b**). In many previous studies, when all Arg residues were mutated to Lys in a sequence with a significant fraction of these residues (13.7% Arg in LAF-1 RGG, 10.2% in DDX4)^19, 24^, no measurable droplet formation was observed^19, 24, 26^; this was interpreted by many that Arg is required for LLPS of PLCDs at physiologically relevant conditions. Additionally, based on the saturation curves for the G<u>K</u>G<u>N</u>SPYS and the G<u>Q</u>G<u>N</u>SPYS variants in **Fig. 3a**, G<u>K</u>G<u>N</u>SPYS may phase separate more avidly than G<u>Q</u>G<u>N</u>SPYS at higher concentrations, highlighting the significance of mapping LLPS at a range of temperatures and concentrations.

To elucidate the origin of the differences in interactions of Lys and Arg with the aromatic group, we computed the potential of mean force between the cationic atoms and the atoms of the Tyr *π*-ring in the GRG<u>N</u>SPYS and G<u>K</u>G<u>N</u>SPYS variants from the atomistic condensed phase simulations. We observe that a free-energy minimum exists at ∼5Å for the Arg:Tyr interactions that is absent for Lys:Tyr (**Fig. S11**). When considering the orientations of the *π*-ring to the sidechain of cationic residues at shorter distances (**Fig. 3g**), a significant fraction of configurations involve stacking of the *π*-ring above or below the Arg sidechain^20, 47^, which is not observed in the case of Lys. Furthermore, when considering contacts formed by Arg or Lys with other residues, we observe that all contacts involving Lys are decreased, whereas other contacts remain relatively unchanged compared to the polycationic Arg variant (**Fig. 3h**). In addition to Tyr, the contacts of Lys with polar residues such as Ser and Asn also show a pronounced reduction^48^, implying that these often ignored interactions of Arg with other residues may be essential in LLPS.

To test the potential role of polar residues in LLPS, we mutated the Asn and Ser residues at positions 4 and 8 of the GRG<u>N</u>SPY<u>S</u> repeat unit to Ala. The phase separation of this new variant GRG<u>A</u>SPY<u>A</u> is measurably reduced, as presented in the comparison of their phase diagrams (**Fig. S12**). This is an unexpected result based on the previous literature and similar substitutions introduced earlier in this paper, but consistent with the elevated role of these interactions in the polycationic sequence.

### 2.5 Polycationic Arg-rich sequence phase separates without any aromatic residue

The A-IDP variants presented to this point each contain aromatic Tyr, which has been shown to be sufficient to induce LLPS in a variety of natural and synthetic proteins^23, 29^. Our simulation data above identified a diverse array of interactions with Tyr, in addition to the commonly expected aromatic-aromatic interactions between Tyr residues. To clearly separate Tyr’s contribution, we carried out Tyr-to-Ala substitutions in the GRG<u>N</u>SP<u>Y</u>S repeat to generate a new variant, (GRG<u>N</u>SP<u>A</u>S)_25_, which lacks most interactions that are currently understood to be essential for LLPS. The comparison between pairwise contacts between the (GRG<u>N</u>SP<u>Y</u>S)_25_ and (GRG<u>N</u>SP<u>A</u>S)_25_ variants highlights the loss of all residue pairwise contacts involving Tyr-to-Ala substitution, and not just self-interactions (**Fig. 4a**, **Fig. S14**); this is further evidence that aromatic Tyr contacts with nonaromatic residues are also essential in LLPS.

**Fig. 4.**
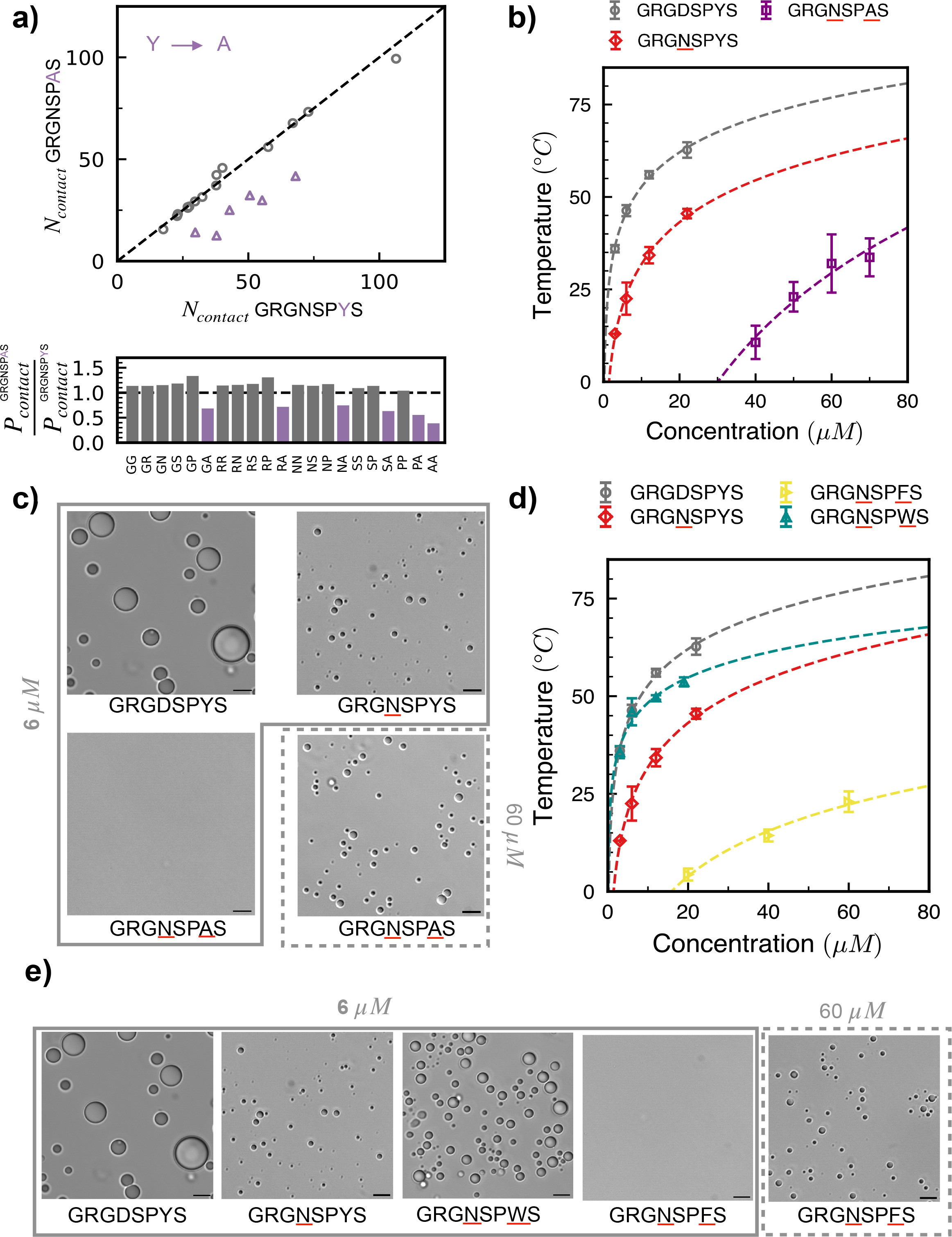
Aromatic residues promote but are not required for phase separation. (a) Average residue pair-wise contacts estimated through atomistic simulations for GRGNSPAS with respect to the GRGNSPYS variant. Residue pairs not involving mutated residues are shown as gray circles while residue pairs involving the mutated residues (Y to A) are shown as purple triangles. The plot below shows a ratio between the probability of contact formation of each of the residue pairwise contacts for the GRGNSPAS variant to that of the GRGNSPYS variant. (b) Partial phase diagrams for WT (GRGDSPYS), GRGNSPYS, and GRGNSPAS in PBS. Dashed lines in the left plot are obtained through a logarithmic fit to the measured transition temperatures at each concentration from turbidity experiments. (c) Microscopy for WT, GRGNSPYS, and GRGNSPAS variants at a concentration of 6 µM, shown in the box bounded by solid lines, and additionally for 60 µM for GRGNSPAS, shown in the box bounded by dashed lines. (Scale bar, 5µm) (d) Partial phase diagrams for GRGNSPFS and GRGNSPWS variants in PBS, with WT and GRGNSPYS variants shown as a reference. Dashed lines in the left plot are obtained through a logarithmic fit to the measured transition temperatures at each concentration from turbidity experiments. (e) Microscopy for the WT, GRGNSPYS, GRGNSPFS, and GRGNSPWS variants at a concentration of 6 µM, shown in the box bounded by solid lines, and additionally for 60 µM for GRGNSPFS, shown in the box bounded by dashed lines. (Scale bar, 5µm). Note: Phase diagrams and microscopy data for WT and GRGNSPYS are the same as shown in Fig. 2; they are repeated here for reference.

Remarkably, our experimental data shows that the new (GRG<u>N</u>SP<u>A</u>S)_25_ polypeptide can undergo LLPS and form liquid-like droplets (**Fig. 4b**, **Fig. 4c)**. At a polypeptide concentration of 40 µM, the T_t_ was 10.6 ± 4.5°C (**Fig. 4b**, **Fig. S13**). For reference, 140 residue-long α-synuclein was found to undergo phase separation at concentrations above 200 µM in the presence of 10% PEG-8000 at physiological salt conditions^49^. This result shows that although Tyr is an important contributor for phase separation, it is not required for phase separation in the context of our A-IDP sequence.

### 2.6 Different aromatic residues contribute towards phase separation as expected (Trp>Tyr>Phe)

Finally, we also studied the phase behavior of two variants involving Tyr-to-Trp and Tyr-to-Phe substitutions to gain insight into their relative role in LLPS. We find that these changes to the aromatic groups can lead to appreciable changes in the transition temperatures, in a similar manner as shown previously^12, 14, 24^. We studied these mutations in the context of the cationic sequence, comparing the cationic repeat unit described above, GRG<u>N</u>SP<u>Y</u>S, to repeat units GRG<u>N</u>SP<u>F</u>S and GRG<u>N</u>SP<u>W</u>S. We observe a measurable transition for the (GRG<u>N</u>SP<u>F</u>S)_25_ variant only above 20 *μ*M, versus 3 *μ*M for the Tyr-variant (**Fig. S13**). The phase diagram of this variant is significantly shifted downward, highlighting reduced LLPS, although the variant still exhibits the ability to form droplets at 60 µM at 20 °C. In contrast, LLPS of the (GRG<u>N</u>SP<u>W</u>S)_25_ variant is significantly enhanced, approaching that of the charge-neutral WT A-IDP, (GRGDSPYS)_25_ (**Fig. 4d**). Microscopy images confirm the liquid-like morphologies of the condensates formed by these variants (**Fig. 4e**). All-atom simulations also corroborate these findings, as shown by the comparison of the pairwise contacts between different variants (**Fig. S14**, **Fig. S15**). Our findings follow the expected trends of Trp>Tyr>Phe, in terms of contribution to LLPS, as reported previously in many other natural and synthetic proteins^12, 14, 24, 26^. Somewhat surprisingly, the phase diagrams of (GRG<u>N</u>SP<u>F</u>S)_25_ and (GRG<u>N</u>SP<u>A</u>S)_25_ overlay approximately, demonstrating a similar contribution of aromatic Phe versus aliphatic Ala residues in the context of the polycationic sequence.

### 2.7 A-IDP variants yield polypeptide sequences with tunable material properties

The material properties that emerge upon LLPS are critical to the function, dysfunction, and engineering of biomolecular condensates. We therefore sought to understand not only how sequence determines phase behavior, but also how amino acid sequence relates to material properties, specifically condensate rheology. Remarkably, all the A-IDP sequences tested maintained spherical droplet morphology over the course of 24 hrs with no observed fibrillization or aggregation (**Fig. S16**). To quantify the rheology of A-IDPs that undergo LLPS, we adapted a passive microrheology technique based on the Brownian motion of fluorescent tracer nanospheres embedded in the condensates^31, 50, 51^. Using this approach, we evaluated the viscosities of the WT and nine variants for which microrheology was experimentally feasible. Tracer beads embedded within these condensates displayed Brownian motion (**Fig. 5a**). The beads’ mean squared displacements (MSDs) increased linearly with lag time *τ* for all the variants, and by fitting MSDs to the power law *MSD*(*τ*) = 4*Dτ*^*α*^, we obtained exponent values close to 1 (**Fig. 5b**), indicating these condensates behave largely as viscous fluids under the given experimental conditions. Using the diffusion coefficient *D* obtained from the MSD data, we calculated condensate viscosity *η* using the Stokes-Einstein relation (see Methods).

**Fig. 5.**
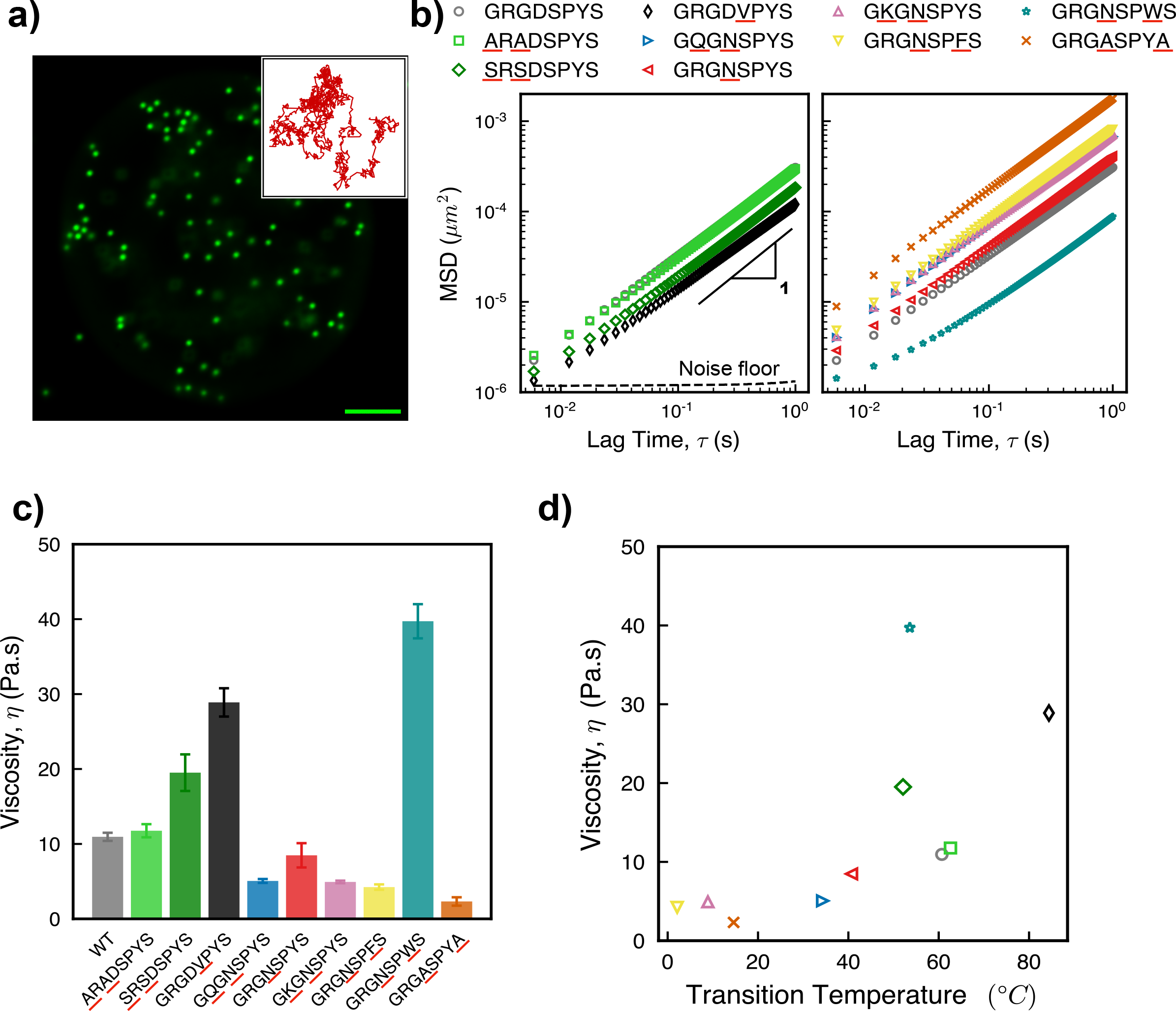
Variants result in condensates with diverse material properties. (a) Fluorescence microscopy image of 0.5 µm yellow-green fluorescent polystyrene beads embedded in WT droplet (Scale bar, 5 µm). Inset: Representative trajectory from two-dimensional particle tracking showing Brownian motion of the beads (length of inset box represents 0.02 µm). (b) Ensemble mean-squared displacement versus lag time for the variants tested in this study. (GQGDSPYS is not shown because it did not phase separate, and GRGNSPAS condensates were too small to analyze using our microrheology method.) (c) Viscosity of the variants, calculated from the particle tracking results. (d) State diagram showing transition temperatures and viscosities for the variants tested in this study. Transition temperatures are calculated at 18 µM concentration, whereas for viscosity measurements, total concentration differed for different variants. The symbols and colors on the plot are in accordance with the legend of subplot (b).

Overall, we find that the viscosity varies by more than one order of magnitude among the sequences examined (**Fig. 5c**). The viscosity of WT was measured as ∼ 11 Pa.s, approximately 11,000 times the viscosity of water. Out of nine variants tested, the lowest measured *η* value was for GRG<u>A</u>SPY<u>A</u>, at ∼ 2 Pa.s. In contrast, the highest measured *η* was for the Trp-containing variant, GRG<u>N</u>SP<u>W</u>S, with *η* ∼ 40 Pa.s. This 20-fold range in viscosity highlights the diverse material properties accessible through mutating IDP sequence and thereby altering molecular interactions.

To explore the relationship between transition temperature (T_t_) and material properties for the polypeptide sequences, we plotted the viscosity vs. transition temperature for the sequences (where T_t_ is calculated at A-IDP concentration of 18 µM, and viscosities were all measured at ambient temperature). Interestingly, we observed only a moderate correlation (**Fig. 5d**) between *η* and T_t_, with the Pearson correlation coefficient determined to be 0.67.

We observe that in most of the neutral and cationic polypeptides (G<u>Q</u>G<u>N</u>SPYS, GRG<u>N</u>SPYS, GRG<u>N</u>SP<u>F</u>S, and GRG<u>A</u>SPY<u>A</u>), *η* is reduced as compared to the zwitterionic WT sequence. This is consistent with the lowering of T_t_ for these variants as compared to WT. A notable exception, however, is the GRG<u>N</u>SP<u>W</u>S variant. A four-fold increase in *η* of GRG<u>N</u>SP<u>W</u>S is observed despite a decreased T_t_ as compared to WT, suggesting that the bulky indole side chain may slow the dynamics of the condensates. We performed fluorescence recovery after photobleaching (FRAP) on WT and GRG<u>N</u>SP<u>W</u>S to verify this microrheology result, and indeed, GRG<u>N</u>SP<u>W</u>S exhibits significantly slower FRAP recovery as compared to WT (**Fig. S17a**).

Further examination of the viscosity data set reveals several interesting and even surprising results. First, we measured only a two-fold reduction in *η* for the R to K mutation (GRG<u>N</u>SPYS vs. G<u>K</u>G<u>N</u>SPYS), a smaller decrease than might be expected based on the 300-fold reduction observed in polyK as compared to polyR complex coacervates with UTP^31^. Second, microrheology of the G to S and G to A variants confirms that Gly is not necessary for fluidity of these A-IDP condensates. We observed that *η* of <u>A</u>R<u>A</u>DSPYS is comparable to that of WT, and the <u>S</u>R<u>S</u>DSPYS sequence likewise remained liquid. However, <u>S</u>R<u>S</u>DSPYS exhibited a two-fold increase in *η* compared to WT despite similar T_t_, which represents another exception to the correlation between transition temperature and viscosity. This result suggests that replacing Gly with Ser enhances the strength of intermolecular interactions, perhaps due to presence of hydrogen bonding that lowers the mobility and dynamics of the proteins. Third, a 3-fold increase in *η* for GRGD<u>V</u>PYS (∼30 Pa.s) as compared to WT suggests that mutating polar Ser to bulky hydrophobic Val promotes enhanced intermolecular hydrophobic interactions within the condensates and results in slowed droplet dynamics and increased viscosity.

Collectively, these observations indicate that a range of viscosities can be obtained by targeting specific amino acid residues and shed light on the sequence-material properties relationship of the polypeptide condensates. These results also suggest that a correlation exists between transition temperature and viscosity; however, the exceptions we observed in this study point to the feasibility of independently modulating phase separation propensity and material properties.

## 3. Discussion

In this work, we show that the phase separation propensity of IDPs is a result of multiple interactions between diverse residue pairs in a sequence; it is not solely dictated by the presence or absence of a limited subset of residues. Based on experiments and simulations of repetitive model polypeptides, we find that polar and nonpolar residues that have not been considered as significant for phase separation propensity in the current literature in fact form a multitude of interactions with all other residues in the sequence, aiding in phase separation. The collective presence of a sufficient number of such interacting residues in an IDP tips the balance towards phase separation^52^. This work therefore points to the diverse range of IDP sequences that can phase separate under biologically relevant conditions. We tested a number of state-of-the-art sequence-based predictors^20, 34–36, 53, 54^ on the A-IDPs used in this study and found that the results did not match the experimental findings detailed in this work, underlining that much of the sequence-phase behavior relationship is still not understood (**Fig. S18**). The expanded molecular language presented in this work, accounting for all residues, suggests opportunities for improved bioinformatics prediction of proteins that phase separate in vivo. Furthermore, our work highlights that a diverse palette of sequences can be constructed to generate protein-based biomaterials with controllable material properties for biotechnology applications.

The observation of phase separation of many of the different polypeptides examined here, despite the substantive changes to the WT sequence, imply that a combination of various interactions involving all residues contribute to the overall phase separation propensity of the sequence (**Fig. 6a**). These diverse interactions – including electrostatic, sp^2^/*π* or *π* − *π*, cation-*π*, hydrogen bonding, and hydrophobic interactions – work in tandem to allow a given polypeptide to undergo LLPS, but removal of one interaction mode still results in LLPS with a lower phase separation propensity. To highlight this additive effect of interactions on phase separation propensity, we perform a thermodynamic analysis analogous to that presented by Bremer et al.^26^ and Ng et al.^55^ to quantify the changes in saturation concentration upon mutation of a particular residue (**Fig. 6b**). The results of this analysis highlight that certain mutations have a more pronounced effect on phase separation propensity; however, the relative changes of these mutations show a significant dependence on the temperature of the system (**Fig. S19**). Moreover, the derived values for changes in saturation concentration are not necessarily universal – they can depend on the reference sequence used (**Table S1**), highlighting the importance of context dependence.

**Fig. 6.**
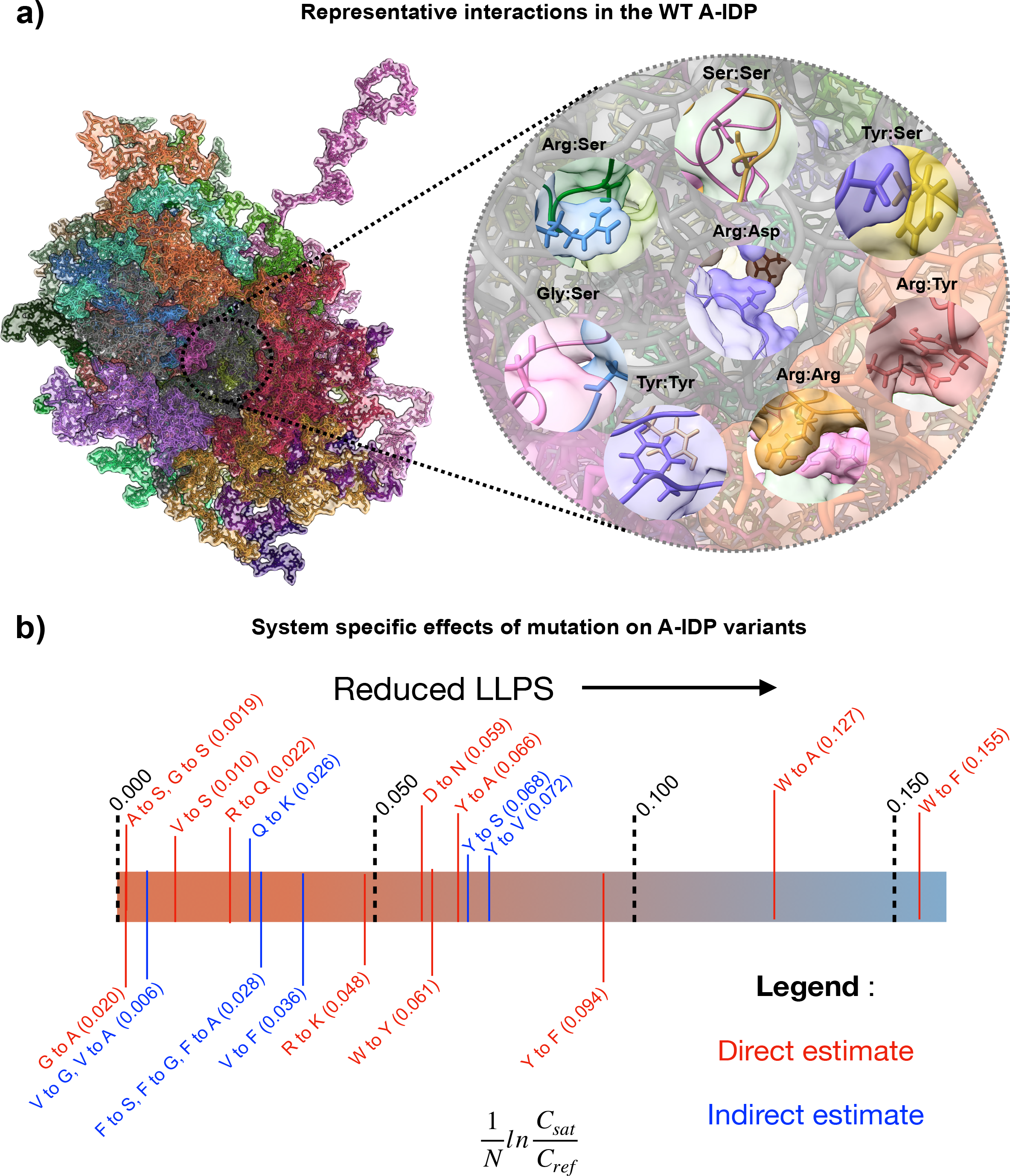
A multitude of interactions work in tandem to drive LLPS. (a) Snapshot from atomistic simulations of WT, highlighting representative examples of the wide variety of contacts driving phase separation of WT. Residues are represented by their three letter amino acid codes. (b) Effect of different mutations to C_sat_ at 37 °C, normalized by the number of mutations carried out (denoted by N). C_sat_ denotes the saturation concentration of the variant, whereas C_ref_ denotes the saturation concentration of the reference sequence used to calculate the effect of the mutation. Direct estimates refer to values for which the particular mutation was carried out in this work, while indirect estimates refer to values where the mutations were not carried out, but can be estimated based on combining data from multiple investigated variants.

It is instructive to consider how our present results compare with earlier work. For instance, several prior studies report the lack of phase separation when Arg residues in the sequence are mutated to Lys^19, 24^, seemingly at odds with the findings presented here, where the GKGNSPYS repeat polypeptide phase separates, with a C_sat_ of ∼25 µM at 20 °C. This discrepancy can be explained by the multiple interactions present in our WT sequence that contribute to LLPS, such that none are essential and LLPS is robust to mutation. Furthermore, a significant portion of previous studies attribute the effect of R to K mutation to the loss of cation-π interactions^18, 19, 21, 26^; the results presented in this work support the hypothesis that the dramatic difference in importance of Arg and Lys cannot be explained by cation-π interactions alone but is a result of reduced interactions between multiple residues as a result of the mutation. Our prior work on LAF-1 RGG demonstrated that aside from electrostatic cation-π interactions, hydrogen bonding between Arg and Tyr contribute to phase separation^24^. Here, we show that the differences between Arg and Lys, though notable for their divergent strength of interactions with aromatic groups as previously shown^45, 46^, also extend to their interactions with all other residues.

Previously, several theories have suggested a correlation between phase separation temperature and viscosity of condensates, and this correlation was shown experimentally for R/G-repeat polypeptide-RNA coacervates^50, 56, 57^. The current study, however, explores whether such a relationship exists for polypeptide sequences devoid of any nucleic acid interactions and links the material properties of these condensates to the sequence. On the one hand, our results suggest that the changes in thermodynamic phase behavior could be a useful proxy for predicting the trends in condensates’ viscosity. On the other hand, these results indicate that phase separation propensity and rheology may not always be coupled. This may be due to various factors, one of which is suggested by the GRGNSPWS sequence: that bulky side chains may increase condensate viscosity.

Another consideration is that thermodynamic interactions that drive phase separation may be less relevant to viscosity than dynamic interactions that emerge once the condensate has formed^58^. Understanding the coupling – or uncoupling – of phase behavior and rheology is vital for understanding how disease-related mutations may perturb rheology, and it is also useful for the aim of generating designer materials. It is important to note that the sequence-to-rheology relationship must be considered in the context of the particular experimental system. The purely viscous behavior of these sequences, as opposed to viscoelastic behavior observed in the polypeptide-nucleic acid sequences shown before^50^, may be due to the role of nucleic acids in the previous study. As another example, Gly to Ala mutation had little effect on rheology of our sequence, which differs from the result for FUS, where G to A mutation significantly slows droplet fusion and causes droplet hardening^21^.

Taken together, the results presented in this work highlight two main points. First, phase separation propensity is a result of multiple interactions between diverse residue pairs. Second, the extent to which each residue contributes to phase separation is dependent on context, such as the temperature and surrounding amino acid sequence (and other contextual factors beyond the scope of the current study, such as sequence patterning). We believe that insights provided by experiments and atomistic simulations regarding the relationship between sequence, phase behavior, and material properties will significantly aid in sequence-based prediction and design of biomolecular condensates.

## 4. Material and methods

### 4.1 Materials

The plasmid DNA encoding the different A-IDP sequences in pQE80L cloning vectors was purchased from Genscript Corporation (Piscataway, NJ). Chemically competent cells of E. coli strain M15-[pREP4] (for transformation of recombinant plasmids) and RNAse (for protein purification) were purchased from Qiagen (Valencia, CA). All other chemicals were obtained from Sigma-Aldrich (St. Louis, MO) or Fisher Scientific (Waltham, MA) and were used as received unless otherwise noted.

### 4.2 Protein Expression and Purification

The E. coli M15-[pREP4] strain was transformed with each DNA plasmid (pQE80L cloning vectors) by heat shock to generate the expression cell stocks employed in protein production. Protein expression and purification were conducted as previously reported by our laboratories_38,59-61_. The complete protocol can be found in the Supplementary Methods section.

### 4.3 General Characterization of A-IDPs

The purity of samples was assessed by SDS-PAGE using NuPAGE 4-12% Bis-Tris gels (Invitrogen) and stained using a Coomassie stain (GelCode Blue Safe Protein Stain; Invitrogen). For this purpose, the lyophilized samples were dissolved in PBS buffer at a concentration of 1 mg/mL, via sonication for 2 min with a 10 s recovery, and diluted with 8M urea buffer (pH 8.0) to a final polypeptide concentration of 0.5 mg/mL (**Fig. S20**). Amino acid analysis was performed by the Molecular Structure Facility at the University of California, Davis (Davis, CA) using a Hitachi L-800 sodium citrate-based amino acid analyzer (Tokyo, Japan) to determine the composition of each polypeptide (**Table S2, Table S3**). The purity and molecular weight of the dialyzed polypeptides (before lyophilization) was confirmed via UPLC and electrospray ionization mass spectrometry (ESI-MS) (Waters Xevo G2-S Q-TOF MS with Acquity UPLC, Milford, MA) (**Fig. S21, Fig. S22**). The A260/A280 ratios were measured in a UV-Vis spectrophotometer (Cary 60 Bio; Agilent), and the resulting ratios for all sequences range between 0.55 – 0.65, which suggests the absence of any remaining nucleic acid.

### 4.4. Turbidity Assays

#### a) Kiick Laboratory

Temperature-dependent turbidity assays were conducted in a UV-Vis spectrophotometer (Cary 60 Bio; Agilent) equipped with a Peltier temperature controller. Protein samples were tested in low-volume quartz cuvettes with 1 cm path length (Hellma Analytics). Lyophilized samples were dissolved in PBS (8 mM Na_2_HPO_4_, 137 mM NaCl, 3mM KCl, 2mM KH_2_PO_4,_ pH 7.4) at double the desired concentration and were sonicated for 2 min with a 10 s recovery time, allowing them to warm up during sonication and exceed the transition temperature. Then, the samples were filtered (0.45 *µ*m, PVDF) at high temperature to eliminate any non-dissolved material, transferred to the quartz cuvette, and incubated at 80 °C in the spectrophotometer. The measured absorbance values (at λ = 280 nm) and the molar extinction coefficients (obtained from the ProtParam tool at web.expasy.org) were used to calculate the polypeptide concentration, and then this information was used to adjust the sample to the desired final concentration and to a volume of 500 *µ*L in PBS. The samples were cooled from 80 °C to 5 °C at a rate of 1 °C min_-1_ and the absorbance was measured at λ = 350 nm every 1 °C throughout the temperature ramp. The transition temperature was defined as the point where absorbance first exceeds 0.03.

#### **b)** Schuster Laboratory

For verification purposes, temperature-dependent turbidity assays were also conducted in the Schuster Lab in a UV-Vis spectrophotometer (Cary 3500; Agilent) equipped with a multicell Peltier temperature controller. Protein samples were tested in a quartz cuvettes with 1 cm path length (ThorLabs) or low-volume quartz cuvettes with 1 cm path length (Hellma Analytics) and sample preparation was conducted as described above in part (a). The samples were filtered using 0.45 *µ*m, PES filters.

### 4.5. Circular Dichroism Spectroscopy

Circular dichroism (CD) spectroscopy (Jasco J-1500 CD spectropolarimeter, Jasco Inc., Easton, MD, USA) was conducted to characterize the secondary structure of the A-IDP sequences. A description of the sample preparation and the resulting spectra can be found in **Fig. S23**. The structural comparison of each sequence can be found on **Table S4**.

### 4.6. Microscopy: Phase Behavior and Droplet Fusion

Each polypeptide sample was prepared at the desired final concentration by diluting the stock solution (prepared as described in section 4.4) with PBS (pH 7.4). Prior to imaging, samples were kept at 70 °C in a heat block (Fisher Scientific) in the microscope room, so that there was no droplet assembly before the start of the experiment. The dishes used for microscopy were 16-well glass-bottom dishes (#1.5 glass thickness; Grace Bio-Labs) that were pretreated with 5% Pluronic F-127 (Sigma-Aldrich) for a minimum of 45 minutes. The coated wells were washed with PBS (pH 7.4), and then 100 *µ*L of the protein sample was removed from the heat block and transferred to the imaging well at room temperature to initiate droplet assembly.

Imaging of droplet assembly at different time points after the plating (considered t = 0) was performed on a Zeiss Axio Observer 7 inverted microscope equipped with an Axiocam 702 monochrome sCMOS camera (Zeiss), employing a 63x/1.4 NA plan-apochromatic oil-immersion objective and using differential interference contrast (DIC) transillumination. The same Zeiss microscope and experimental conditions were used for observing droplet fusion, with imaging conducted at a frame rate of approximately 200 Hz. Videos of droplet fusion events were analyzed using MATLAB. All microscopy experiments were conducted at the ambient temperature (17-20 °C).

### 4.7. Video Particle-Tracking Microrheology (VPT)

500 nm diameter yellow-green carboxylate-modified polystyrene beads (FluoSpheres, Invitrogen) were used for VPT microrheology measurements. Each polypeptide sample was prepared at the desired final concentration for microrheology by diluting the stock solution (prepared as described in sections 4.4 and 4.6) with PBS (pH 7.4). To employ the microrheology technique for all the mutants, which have a wide range of saturation concentrations, we used higher polypeptide concentrations for the variants with elevated saturation concentrations (G<u>K</u>G<u>N</u>SPYS, GRG<u>N</u>SP<u>F</u>S, and GRG<u>A</u>SPY<u>A</u>). Since viscosity is an intrinsic material property of the condensate, it should not change with total polypeptide concentration. We verified this experimentally for two polypeptide sequences (**Fig. S17b**). This permits us to compare the viscosities of the variants, even though polypeptide concentration varied for the different samples.

Microrheology experiments were prepared by mixing the 200 µL polypeptide sample with the fluorescent tracer beads before initiating droplet assembly in a 96-well plate (#1.5 high-performance cover glass, Cellvis). The samples were incubated at room temperature for 45 min and then were observed under the microscope to verify that the tracer beads were embedded in the condensates (**see Fig 5a, main text**). Next, the samples in the well plate were centrifuged at 300xg for 1 minute to form a condensate layer or larger-size droplets (>30 *µ*m in diameter); the purpose of this step was to avoid boundary effects and prevent flow of the condensates.

Epifluorescence video imaging was initiated at the 1 hr timepoint using the same microscope and procedure as described in section 4.6, with fluorescence excitation using a 475 nm LED (Colibri 7; Zeiss). Videos of the tracer beads diffusing within the condensate were collected at 200 frames per second for 2000 frames. Imaging was conducted at room temperature (17-20 °C). For each A-IDP variant, two independent samples were made on different days, and 4-5 videos were collected from each sample, with each video containing ∼10-50 tracer beads.

The TrackPy particle tracking code (see Section 4.8) was used to analyze the collected videos, starting with extracting particle trajectories. The mean squared displacement (MSD) was calculated from the trajectories of individual beads, followed by calculating the ensemble-average MSD. In general, the ensemble-average MSD often scales as a power law with lag time *τ*, as given by the following equation:

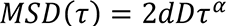

where d is the number of dimensions (here d=2, since data collection and analysis were conducted in the x-y plane), D is the diffusion coefficient, and α is the diffusivity exponent. For a purely viscous fluid, the diffusivity exponent α is close to unity. α values for all the condensates tested were in the range of 0.82-1.05. Assuming a purely viscous fluid, with the system at equilibrium, the condensate viscosity η is then calculated using the Stokes-Einstein equation:

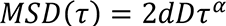

where k_B_ is the Boltzmann constant, T is the temperature (Kelvin), and R is the tracer bead radius. Reported viscosities (**Table S5**) are averages ± SD across multiple videos.

The noise floor of 500 nm beads was measured by adhering the beads to the glass surface of a 96-well plate. We acquired the trajectories of the beads adhered to the glass surface using the same parameters as those used for VPT studies of the polypeptide samples. We find that the particles adhered on the glass surface showed negligible change in MSD over lag time, with MSD measured at approximately 1.1×10_-6_ *µ*m^2^ (Fig. 5b).

### 4.8. Software

VPT data analysis was conducted using the open-source particle tracking package TrackPy (v0.5.0)_62_ in Python and customized as needed. Custom MATLAB code_24_ was used to analyze droplet fusion events. Fiji (version 1.53s) was also used for image processing.

### 4.9. All-Atom Molecular Dynamics simulations

#### 4.9.1 Generating and running the atomistic WT slab

The initial configuration for the all-atom slab was generated from a Coarse-Grained (CG) simulation using the steps detailed in prior work^39^. The initial CG simulation consisted of 30 chains of the WT peptide using the HPS-Urry model^63^, run for 1 microsecond. The final configuration of the CG simulation was backmapped into an atomistic representation using MODELLER^64^. Following this, the atomistic slab was minimized using CAMPARI^65^ with the implicit solvent ABSINTH^66^ force field. The force field used was Amber99SBws-STQ^67^. This minimized structure was then solvated with TIP4P/2005^68^ water, along with scaled salt interactions^69^, in a box of size 10×10×40 nm^3^. Following solvation, the system was minimized using GROMACS-2019.4 using the steepest descent algorithm. After minimization, 100 mM of NaCl was added to the system in excess of the amount of ions needed to maintain electroneutrality. The solvated polypeptide with ions was then minimized once again using the same parameters as prior minimization and then 100 ns of NVT was run with all bonds constrained using the Nose-Hoover thermostat^70^ with a coupling constant of 1 ps for protein, water, and ions and temperature fixed at 300 K. Following the NVT equilibration, 100 ns of NPT equilibration was run using the Berendsen barostat^71^ with isotropic coupling and a constant of 5 ps for pressure control and the Nose-Hoover thermostat with same parameters as for the NVT run. Following the equilibration steps, a 1 microsecond production run in the NPT ensemble using the Langevin Middle Integrator^72^ and the Monte Carlo Membrane Barostat in OpenMM-7.7^73^ was run. Short-range nonbonded interactions were calculated with a cutoff radius of 0.9 nm, while long-range electrostatics were treated with the PME method^74^. Hydrogen mass was increased by 1.5 times, allowing for a timestep of 4 fs, while hydrogen containing bonds were constrained using the SHAKE algorithm^75^. A friction coefficient of 1 ps^-1^ was used for the Langevin Middle integrator. The membrane barostat was set to be isotropic in x and y dimensions with a free z dimension. Pressure was fixed at 1 bar, temperature at 300K, default surface tension of 200 bar·nm, and a frequency of 1000 time steps for attempted volume changes.

#### 4.9.2 Generation of mutated slabs

A mutant slab was generated from a parent slab using the following steps. The final frame of the parent slab after the production run was separated into two PDB files, one of protein and the other containing water and ions. The protein-only PDB was passed to UCSF Chimera,^76^ where the mutations were carried out. The mutated PDB was then saved and combined with the water+ions PDB from the parent slab. Following the combination of the two PDBs, the system was minimized and equilibrated using the same steps and parameters as the WT detailed above, before a production run of 1 microsecond with the same parameters as WT was run in OpenMM-7.7. In some cases, where the mutation being carried out involves a significant change in the size of the side chain, e.g. the Y to W mutation, a soft-core minimization with lambda set to 0.01 and alpha set to 4 was carried out immediately after combination of the PDBs so as to remove any clashes resulting from protein and water or ion atoms overlapping in the initial PDB. In cases where net charge of the peptide was non-zero, counterions were added to maintain neutrality prior to minimization and equilibration steps. Following this soft-core minimization, the steps remained the same. A flowchart indicating the sequential mutations carried out to generate the set of variants is available in the Supporting Information (**Fig. S24**).

#### 4.9.3 Analysis of all-atom simulation trajectories

Two residues were counted as a contact if any two heavy atoms of the residues are within 6 Å of each other. All analysis shown in the text is calculated after 250 ns of simulation time. The equilibration time of 250 ns was estimated using the autocorrelation of the radius of gyration of the chains in the system calculated for the WT (**Fig. S25**). Inter- and intrachain contacts are calculated together. Details regarding the normalization to generate residue pairwise contacts can be found in work by Zheng et. al.^39^ Calculations of pairwise non-bonded energies are performed using pairwise option in AmberTools21^77^. All snapshots were generated using VMD-1.9.3 and UCSF ChimeraX^78^.

## Acknowledgements

We acknowledge the following grants: NSF DMR-2004796 to KLK and JM. NIH R01GM136917 to JM. Welch Foundation A-2113-202203311 to JM. NIH R35GM142903 to BSS.

## Author contributions

SR, CGG, and MB wrote the draft. All authors edited the draft. CGG expressed and purified all polypeptides and performed turbidity experiments. MB performed all the microscopy and microrheology experiments and conducted additional turbidity experiments. SR performed the simulations. SR and AR analyzed the simulations. BSS, KLK, and JM designed and supervised the research.

