## Supplementary Information for "Expanding the molecular language of protein liquid-liquid phase separation"

**Supplementary Methods:**

#### Protein Expression and Purification

A single colony of *E. coli* M15[pREP4] containing the desired construct was inoculated in 150 mL of sterile LB media containing 100 μg $mL^{-1}$ antibiotics (ampicillin) and grown overnight. Overnight culture media (150 mL) was used to evenly inoculate 6x750 mL of 2xYT media (yeast extract 10 g $L^{-1}$, NaCl 5 g $L^{-1}$, and tryptone 16 g $L^{-1}$) for protein expression. The 750 mL cultures were grown in a shaker at 37°C until the OD600 reached 0.6–0.8, and then isopropyl β-D-1-thiogalactopyranoside (IPTG) was added to a final concentration of 1 mM to induce protein expression. After 8 h of culture for protein expression, cells were harvested by centrifugation (5000 rpm for 15 min at 4°C), and the cell pellets were frozen with liquid nitrogen and transferred to a clean plastic bottle. The frozen cell pellets were thawed and resuspended in pH 8.0 native lysis buffer (50 mM $NaH_2PO_4$, 300 mM NaCl, and 10 mM imidazole) with 0.45 g of lysozyme. Lysed cells were further disrupted via sonication on ice, using a Fisher Scientific model 500 Sonic Dismembrator (10 mm tapered horn) for 20 min with a 10-s recovery time and subsequently incubated with RNAse (10 μg $mL^{-1}$) and DNAse (5 μg $mL^{-1}$) for 30 minutes. The supernatant from centrifugation (20,000 rpm for 15 min at 4°C) of cell lysate was separated and the cell pellet was resuspended in denaturing lysis buffer B (8M urea, 100 mM $NaH_2PO_4$, 10 mM Tris·Cl, pH 8.0) via sonication for 1 min with a 10-s recovery time. The supernatant from centrifugation (20,000 rpm for 15 min at 4°C) was collected and the pH was adjusted to 8.0, followed by incubation with Ni-NTA resin for 1 hour at room temperature. The protein-loaded resin was then loaded into a gravitational flow column, washed with denaturing lysis buffer B, denaturing wash buffer C (8M urea, 100 mM $NaH_2PO_4$, 10 mM Tris·Cl, pH 6.3), denaturing elution buffer D (8M urea, 100 mM $NaH_2PO_4$, 10 mM Tris·Cl, pH 5.9), and finally eluted with 75 mL denaturing elution buffer E (8M urea, 100 mM $NaH_2PO_4$, 10 mM Tris·Cl, pH 4.5). Elution E fractions were carefully transferred and dialyzed (MWCO 3.5 kDa) against deionized water (5 L) at room temperature with at least 7 changes of water before lyophilization. The purified protein yield was approximately 30–50 mg per liter of cell culture.

#### Fluorescence Recovery After Photobleaching (FRAP)

FRAP experiments were performed on a Zeiss Axio Observer 7 inverted microscope equipped with an LSM900 laser scanning confocal module and a 63x/1.4 NA plan-apochromatic oil-immersion objective. WT and $(GRG\underline{N}SP\underline{W}S)_{25}$ were mixed with 5% of RGG-GFP-RGG, which partitions into the condensates and serves as a FRAP probe (here, RGG denotes LAF-1 RGG domain)[1]. A region of approximate radius R = 1.5 μm, within droplets whose radii were approximately 2.5R, was bleached with a 405-nm laser. Subsequent fluorescence recovery of the bleached area was recorded with a 488-nm laser for 1 min. Raw FRAP data was normalized and averaged (n = 4 to 9 separate droplets) to obtain the final FRAP recovery curve.

**DNA sequences between BamHI and HindIII cloning sites in pQE80L cloning vectors. Optimized by Genscript Corporation.**

#### GRGDSPYS (WT)

AGATCTGAGAATCTGTATTTTCAAGGTAGATCTGAATTCAATGGCGGTCGTGGTGATAGCCC
GTATAGCGGTCGTGGTGATAGCCCGTACAGCGGTCGTGGCGACAGCCCGTACAGCGGCCGT
GGCGATTCTCCTTATAGCGGCCGTGGCGACAGCCCGTATAGCGGCCGTGGCGATAGTCCTTA
TTCTGGCCGTGGCGACTCTCCTTATTCTGGACGTGGCGATTCTCCTTACTCTGGCCGTGGCGA
CTCCCCGTACTCTGGTCGTGGCGATTCTCCGTATTCTGGACGTGGCGACTCTCCCTATTCTGG
CCGTGGCGATTCTCCCTATTCTGGGCGTGGCGACTCTCCATATTCTGGGCGTGGCGATTCCCC
TTATAGCGGCCGCGGAGACAGCCCGTACAGTGGCCGTGGCGATAGTCCCTATTCTGGTCGTG
GCGACTCTCCGTATTCTGGTCGTGGCGATTCCCCGTACAGTGGACGTGGCGACAGTCCTTATT
CTGGTCGTGGCGATAGTCCGTATAGTGGCCGTGGCGACAGTCCTTACAGCGGCCGTGGTGAT
TCTCCTTACAGTGGCCGTGGTGACAGCCCGTACAGCGGTCGTGGCGATAGCCCGTATAGCGG
CCGCGGCGATAGCCCGTACAGCGGTACCTAA

#### ARADSPYS

AGATCTGAGAATCTGTATTTCCAGGGTAGATCTGAATTCAATGGTGCGCGTGCGGATAGCCC
GTATAGCGCGCGTGCGGATAGCCCGTACAGCGCGCGTGCGGACAGCCCGTACAGCGCGCGC
GCGGATAGCCCGTACTCTGCTCGCGCGGATAGCCCGTACAGTGCTCGTGCGGATAGCCCGTA
CAGCGCTCGCGCTGATTCTCCGTACTCTGCTCGTGCGGACAGCCCGTATAGCGCGCGCGCTG
ATAGCCCGTACTCTGCCCGAGCTGATAGCCCGTACAGTGCCCGAGCCGATAGCCCGTACTCT
GCACGAGCTGACTCTCCTTATAGCGCGCGCGCTGACTCTCCGTACTCTGCCCGAGCGGACTC
TCCTTATAGCGCTCGAGCAGACTCTCCTTACTCTGCTCGTGCTGACTCTCCCTATAGCGCGCG
CGCTGACAGTCCGTACTCTGCCCGGGCTGATAGCCCGTACAGCGCCCGCGCTGATTCTCCGT
ATAGCGCGCGCGCCGACTCTCCGTACTCTGCACGAGCGGATTCTCCGTACTCTGCCAGAGCC
GATAGCCCGTACAGTGCACGTGCGGATAGCCCGTACAGCGCGCGTGCGGATAGCCCGTATA
GCGCTCGAGCAGATAGCCCGTACAGCGGTACCTAA

#### SRSDSPYS

AGATCTGAGAATCTGTATTTTCAAGGTAGATCTGAATTCAATGGTAGCCGCAGCGATAGCCC
GTATAGCAGCCGCAGCGATAGCCCGTACAGCAGCCGTAGCGACAGCCCGTATTCTTCTCGTA
GCGACAGCCCGTACTCTTCTCGCTCTGATTCTCCGTACAGCAGCCGCAGCGACAGCCCGTAT
AGTTCTCGTTCTGATTCTCCGTATTCTAGTCGCTCTGATAGTCCGTATTCTAGTCGTTCTGATA
GTCCGTATAGTTCTCGCTCTGATAGCCCTTATTCTTCTCGCAGCGACAGCCCGTACAGTTCTC
GCAGTGACAGCCCGTACTCTAGTCGTTCTGATAGCCCTTATAGTTCTCGTAGTGATTCTCCGT
ATAGCTCTAGAAGCGACAGCCCGTATAGCTCTCGCTCTGATAGCCCCTATTCTAGTCGCAGT
GATTCTCCTTACTCTAGTCGCTCTGATAGCCCATATTCTAGTCGCAGCGATTCTCCTTACAGT
TCTCGCAGTGATTCTCCCTATTCTAGTCGCAGCGATAGTCCTTACAGTAGTCGCAGTGATAGT
CCGTATAGTAGTCGCAGCGATAGCCCGTATAGCAGCCGTAGCGATAGCCCGTACAGCTCTAG
AAGCGATAGCCCGTATAGCGGTACCTAA

#### GRGDVPYS

AGATCTGAGAATCTGTATTTCCAGGGTAGATCTGAATTCAATGGCGGTCGTGGCGATGTGCC
GTATAGCGGTCGTGGCGATGTGCCGTACAGCGGTCGTGGCGACGTTCCGTACAGCGGCCGTG
GCGATGTGCCGTACTCTGGCCGTGGCGACGTTCCGTATAGCGGCCGTGGCGATGTCCCTTAT
AGCGGTCGTGGCGACGTGCCGTACTCTGGTCGTGGCGATGTTCCTTACAGCGGCCGTGGCGA
CGTGCCTTACTCTGGCCGTGGCGATGTCCCCTATTCTGGCCGTGGCGACGTCCCGTACAGTG
GTCGTGGCGATGTTCCATATAGTGGCCGTGGCGACGTTCCATACAGCGGTCGTGGCGATGTG
CCATATAGCGGCCGCGGAGACGTTCCGTACAGCGGACGTGGCGATGTTCCCTATTCTGGACG

TGGCGACGTGCCTTATTCTGGCCGTGGCGATGTACCTTATAGCGGCCGTGGCGACGTCCCTT
ACTCCGGCCGTGGCGATGTACCCTATAGCGGCCGTGGCGACGTACCGTATAGCGGTCGTGGT
GATGTTCCGTACAGCGGTCGTGGTGACGTGCCGTACAGCGGTCGTGGCGATGTTCCGTACAG
CGGCCGCGGCGATGTTCCGTATAGCGGTACCTAA

**GQGNSPYS**
AGATCTGAGAACCTGTACTTCCAGGGTAGATCTGAATTCGATCCGGGTCAGGGTAACAGCCC
GTACAGCGGTCAAGGCAACAGCCCGTATAGCGGTCAGGGCAATAGCCCGTATAGCGGCCAG
GGTAATAGCCCGTACAGCGGCCAGGGCAATAGCCCGTACAGCGGTCAGGGCAACAGCCCGT
ACAGCGGCCAAGGTAATAGCCCGTATAGCGGTCAAGGTAATTCTCCTTACAGCGGCCAAGG
TAACAGCCCGTATAGCGGCCAAGGAAATTCTCCTTACTCTGGTCAAGGAAATTCTCCGTATA
GCGGCCAGGGAAATAGTCCTTATAGCGGCCAGGGAAACTCTCCGTATAGCGGCCAAGGCAA
TAGCCCGTATTCCGGACAAGGTAACTCTCCTTATAGCGGCCAAGGAAACTCTCCTTACTCTG
GCCAAGGCAACTCTCCTTATAGCGGTCAGGGAAATAGTCCTTACTCTGGTCAGGGTAATTCT
CCTTATAGCGGTCAAGGTAACTCTCCTTACAGCGGCCAGGGAAATAGCCCTTATTCTGGCCA
AGGAAATAGCCCTTACTCTGGCCAGGGCAATTCTCCTTACAGCGGTCAGGGTAATTCTCCCT
ATTCCGGACAGGGCAATTCTCCTTACTCTGGTACCTAA

**GQGDSPYS**
AGATCTGAGAACCTGTACTTCCAGGGTAGATCTGAATTCGATCCGGGTCAGGGTGATAGCCC
GTACAGCGGTCAAGGCGACAGCCCGTATAGCGGTCAGGGCGACAGCCCGTACAGCGGCCAG
GGTGATAGCCCGTATAGCGGCCAAGGCGATTCTCCTTATAGCGGTCAGGGTGATTCTCCTTA
CAGCGGCCAGGGCGATTCTCCCTATAGCGGCCAGGGTGACTCTCCTTACAGCGGTCAAGGCG
ATTCTCCATATAGCGGCCAGGGCGATAGCCCGTATAGCGGTCAAGGTGACAGCCCGTATTCT
GGCCAGGGTGACTCTCCCTACAGCGGCCAAGGTGACAGCCCGTACAGCGGACAGGGTGATT
CTCCGTATTCCGGACAAGGTGATTCTCCTTATAGTGGCCAGGGCGATTCTCCATACAGCGGT
CAAGGTGATTCTCCCTATAGCGGTCAAGGAGATTCTCCGTACAGCGGTCAGGGAGACAGCCC
GTATAGTGGTCAGGGTGACTCTCCATACAGCGGTCAGGGAGATTCTCCGTATAGCGGTCAGG
GTGACAGCCCGTATAGTGGCCAAGGAGACTCTCCGTATTCTGGCCAGGGCGATAGTCCTTAT
TCCGGACAGGGAGATAGTCCGTATAGCGGTACCTAA

**GRGNSPYS**
AGATCTGAGAACCTGTACTTCCAGGGTAGATCTGAATTCGATCCGGGTCGTGGTAACAGCCC
GTACAGCGGTCGTGGCAACAGCCCGTATAGCGGCCGCGGCAATAGCCCGTATAGCGGCCGT
GGTAACAGCCCGTATAGCGGTCGCGGTAATAGCCCGTATAGCGGTCGTGGTAATAGCCCGTA
CAGCGGCCGCGGCAACAGCCCGTACAGCGGCCGTGGCAATAGCCCGTACAGCGGTCGCGGA
AATTCTCCTTACAGCGGTCGCGGAAACTCTCCTTACTCTGGTCGAGGCAATTCTCCTTATAGC
GGCCGCGGTAATTCTCCTTACAGCGGCCGTGGAAATTCTCCCTACAGCGGTCGCGGCAACAG
CCCGTACTCCGGACGTGGAAATTCTCCATATAGCGGTCGTGGTAACTCTCCTTATAGCGGCC
GTGGAAACTCTCCGTATTCCGGTCGTGGAAACAGTCCTTATAGCGGGCGCGGTAATTCTCCC
TACAGCGGCCGTGGGAACTCTCCATACTCTGGTCGTGGAAATTCTCCGTATTCTGGCCGCGG
TAATTCTCCATACAGCGGCCGCGGTAATAGTCCTTATAGCGGTCGTGGCAATAGCCCGTATT
CCGGACGCGGCAACTCTCCTTATAGCGGTACCTAA

**GRGNSPWS**
AGATCTGAGAACCTGTACTTCCAGGGTAGATCTGAATTCGATCCGGGTCGTGGTAACAGCCC
GTGGAGCGGCCGCGGCAATAGCCCGTGGAGCGGCCGTGGCAACAGCCCGTGGAGCGGTCGC
GGCAACTCTCCTTGGAGCGGCCGCGGAAATAGCCCGTGGAGCGGTCGTGGCAATTCTCCTTG
GAGCGGCCGTGGTAATTCTCCTTGGAGCGGTCGCGGAAACTCTCCGTGGAGCGGCCGCGGTA
ATTCTCCCTGGAGCGGTCGCGGAAACAGTCCGTGGAGCGGCCGAGGCAACTCTCCCTGGAG

CGGCCGCGGAAACAGCCCTTGGAGCGGTCGCGGCAACAGTCCTTGGAGCGGTCGAGGCAAT
AGCCCGTGGTCCGGACGCGGCAACTCTCCATGGAGCGGCCGCGGGAACTCTCCATGGAGCG
GTCGCGGTAATTCTCCGTGGTCTGGTCGCGGAAACAGCCCCTGGAGCGGCCGCGGGAACTCT
CCGTGGAGCGGTCGCGGGAACTCTCCGTGGAGTGGTCGCGGGAATAGTCCTTGGTCTGGCCG
CGGCAACTCTCCCTGGTCTGGCCGCGGAAACAGCCCATGGAGCGGCCGCGGTAACAGCCCG
TGGTCCGGACGTGGTAATAGCCCGTGGAGCGGTACCTAA

**GRGNSPFS**
AGATCTGAGAACCTGTACTTCCAGGGTAGATCTGAATTCGATCCGGGTCGTGGTAACAGCCC
GTTCAGCGGTCGTGGCAACAGCCCGTTTAGCGGCCGCGGCAATAGCCCGTTTAGCGGCCGTG
GTAACAGCCCGTTTAGCGGTCGCGGTAATAGCCCGTTTAGCGGTCGTGGTAATAGCCCGTTC
AGCGGCCGCGGCAACAGCCCGTTCAGCGGCCGTGGCAATAGCCCGTTCAGCGGTCGCGGAA
ATTCTCCTTTCAGCGGTCGCGGAAACTCTCCTTTCTCTGGTCGAGGCAATTCTCCTTTTAGCG
GCCGCGGTAATTCTCCTTTCAGCGGCCGTGGAAATTCTCCCTTCAGCGGTCGCGGCAACAGC
CCGTTCTCCGGACGTGGAAATTCTCCATTTAGCGGTCGTGGTAACTCTCCTTTTAGCGGCCGT
GGAAACTCTCCGTTTTCCGGTCGTGGAAACAGTCCTTTTAGCGGGCGCGGTAATTCTCCCTTC
AGCGGCCGTGGGAACTCTCCATTCTCTGGTCGTGGAAATTCTCCGTTTTCTGGCCGCGGTAAT
TCTCCATTCAGCGGCCGCGGTAATAGTCCTTTTAGCGGTCGTGGCAATAGCCCGTTTTCCGGA
CGCGGCAACTCTCCTTTTAGCGGTACCTAA

**GRGNSPAS**
AGATCTGAGAACCTGTACTTCCAGGGTAGATCTGAATTCGATCCGGGTCGTGGTAACAGCCC
GGCGAGCGGCCGCGGCAATAGCCCGGCGAGCGGCCGTGGCAACAGCCCGGCGAGCGGTCGC
GGCAACTCTCCTGCGAGCGGCCGCGGAAATAGCCCGGCGAGCGGTCGTGGCAATTCTCCTGC
GAGCGGCCGTGGTAATTCTCCTGCGAGCGGTCGCGGAAACTCTCCGGCGAGCGGCCGCGGT
AATTCTCCCGCGAGCGGTCGCGGAAACAGTCCGGCGAGCGGCCGAGGCAACTCTCCCGCGA
GCGGCCGCGGAAACAGCCCTGCGAGCGGTCGCGGCAACAGTCCTGCGAGCGGTCGAGGCAA
TAGCCCGGCGTCCGGACGCGGCAACTCTCCAGCGAGCGGCCGCGGGAACTCTCCAGCGAGC
GGTCGCGGTAATTCTCCGGCTAGCGGTCGCGGAAACAGCCCCGCGAGCGGCCGCGGGAACT
CTCCGGCGAGCGGTCGCGGGAACTCTCCGGCCAGCGGTCGCGGGAATAGTCCTGCTTCTGGC
CGCGGCAACTCTCCCGCTAGCGGCCGCGGGAATAGTCCTGCCTCTGGCCGTGGTAATTCTCC
CGCGTCCGGACGCGGCAACAGTCCCGCGAGCGGTACCTAA

**GKGNSPYS**
AGATCTGAGAACCTGTACTTCCAGGGTAGATCTGAATTCGACCCGGGCAAGGGCAACAGCC
CGTACAGCGGTAAAGGCAACAGCCCGTATAGCGGCAAAGGTAATAGCCCGTATAGCGGTAA
GGGTAACAGCCCGTACAGCGGCAAAGGTAACAGCCCGTATAGCGGTAAAGGTAATAGCCCG
TACAGCGGCAAGGGTAATTCTCCTTATAGCGGCAAAGGCAATTCTCCTTACAGCGGCAAAGG
CAATAGCCCGTACAGCGGTAAGGGAAACTCTCCTTACAGCGGCAAGGGTAACTCTCCTTATT
CTGGCAAGGGAAATTCTCCCTACTCTGGCAAAGGTAATTCTCCTTACAGCGGTAAGGGCAAT
AGCCCGTACTCCGGAAAAGGTAACTCTCCTTATAGTGGCAAGGGAAATAGTCCTTACTCTGG
CAAGGGAAATAGCCCTTATTCTGGCAAAGGAAATTCTCCCTACAGTGGCAAAGGTAATAGTC
CTTATTCTGGAAAGGGAAACTCTCCCTACAGCGGCAAAGGTAACTCTCCTTACTCTGGAAAA
GGTAATTCTCCTTACTCTGGCAAGGGTAATTCTCCCTACAGCGGTAAAGGTAACAGCCCGTA
CTCCGGAAAGGGAAATTCTCCTTATAGCGGTACCTAA

**GRGASPYA**
AGATCTGAGAATCTGTATTTTCAGGGTAGATCTGAATTCGATCCGGGTCGTGGTGCGAGCCC
GTATGCGGGTCGTGGTGCGAGCCCGTACGCGGGTCGTGGCGCGAGCCCGTACGCGGGCCGT
GGCGCGAGCCCGTATGCTGGCCGTGGCGCGTCTCCGTATGCCGGTCGTGGCGCGTCTCCTTA

TGCCGGCCGTGGCGCGTCTCCTTACGCTGGTCGTGGCGCGTCTCCCTATGCAGGCCGTGGCG
CGTCTCCCTACGCTGGACGTGGCGCGTCCCCTTATGCGGGACGTGGCGCGTCTCCGTACGCC
GGCCGTGGCGCGAGTCCTTATGCGGGCCGTGGCGCGTCTCCATATGCGGGTCGTGGCGCTAG
CCCGTACGCCGGACGTGGCGCGAGTCCTTACGCTGGGCGTGGCGCGTCCCCCTATGCTGGAC
GTGGCGCGAGTCCCTATGCTGGGCGTGGCGCGTCACCTTATGCTGGTCGTGGCGCGTCCCCG
TACGCCGGTCGTGGCGCGAGTCCTTACGCAGGACGTGGCGCGTCCCCGTATGCGGGCCGTGG
TGCGAGCCCGTACGCTGGCCGTGGTGCGTCTCCTTACGCGGGTCGTGGTGCTAGCCCGTATG
CGGGACGTGGTGCGAGCCCGTATGCGGGTACCTAA

**Complete peptide sequences expressed in *E. coli* (including His tag).**

**GRGDSPYS (WT)**
MRGSHHHHHHGSRSENLYFQGRSEFNGGRGDSPYSGRGDSPYSGRGDSPYSGRGDSPYSGRGDS
PYSGRGDSPYSGRGDSPYSGRGDSPYSGRGDSPYSGRGDSPYSGRGDSPYSGRGDSPYSGRGDSP
YSGRGDSPYSGRGDSPYSGRGDSPYSGRGDSPYSGRGDSPYSGRGDSPYSGRGDSPYSGRGDSPY
SGRGDSPYSGRGDSPYSGRGDSPYSGRGDSPYSGT-

**ARADSPYS**
MRGSHHHHHHGSRSENLYFQGRSEFNGARADSPYSARADSPYSARADSPYSARADSPYSARADS
PYSARADSPYSARADSPYSARADSPYSARADSPYSARADSPYSARADSPYSARADSPYSARADSP
YSARADSPYSARADSPYSARADSPYSARADSPYSARADSPYSARADSPYSARADSPYSARADSPY
SARADSPYSARADSPYSARADSPYSARADSPYSGT-

**SRSDSPYS**
MRGSHHHHHHGSRSENLYFQGRSEFNGSRSDSPYSSRSDSPYSSRSDSPYSSRSDSPYSSRSDSPY
SSRSDSPYSSRSDSPYSSRSDSPYSSRSDSPYSSRSDSPYSSRSDSPYSSRSDSPYSSRSDSPYSSRSD
SPYSSRSDSPYSSRSDSPYSSRSDSPYSSRSDSPYSSRSDSPYSSRSDSPYSSRSDSPYSSRSDSPYSS
RSDSPYSSRSDSPYSSRSDSPYSGT-

**GRGDVPYS**
MRGSHHHHHHGSRSENLYFQGRSEFNGGRGDVPYSGRGDVPYSGRGDVPYSGRGDVPYSGRG
DVPYSGRGDVPYSGRGDVPYSGRGDVPYSGRGDVPYSGRGDVPYSGRGDVPYSGRGDVPYSGR
GDVPYSGRGDVPYSGRGDVPYSGRGDVPYSGRGDVPYSGRGDVPYSGRGDVPYSGRGDVPYSG
RGDVPYSGRGDVPYSGRGDVPYSGRGDVPYSGRGDVPYSGT-

**GQGNSPYS**
MRGSHHHHHHGSRSENLYFQGRSEFDPGQGNSPYSGQGNSPYSGQGNSPYSGQGNSPYSGQGN
SPYSGQGNSPYSGQGNSPYSGQGNSPYSGQGNSPYSGQGNSPYSGQGNSPYSGQGNSPYSGQGN
SPYSGQGNSPYSGQGNSPYSGQGNSPYSGQGNSPYSGQGNSPYSGQGNSPYSGQGNSPYSGQGN
SPYSGQGNSPYSGQGNSPYSGQGNSPYSGQGNSPYSGT-

**GQGDSPYS**
MRGSHHHHHHGSRSENLYFQGRSEFDPGQGDSPYSGQGDSPYSGQGDSPYSGQGDSPYSGQGD
SPYSGQGDSPYSGQGDSPYSGQGDSPYSGQGDSPYSGQGDSPYSGQGDSPYSGQGDSPYSGQGD
SPYSGQGDSPYSGQGDSPYSGQGDSPYSGQGDSPYSGQGDSPYSGQGDSPYSGQGDSPYSGQGD
SPYSGQGDSPYSGQGDSPYSGQGDSPYSGQGDSPYSGT-

**GRGNSPYS**

MRGSHHHHHHGSRSENLYFQGRSEFDPGRGNSPYSGRGNSPYSGRGNSPYSGRGNSPYSGRGNS
PYSGRGNSPYSGRGNSPYSGRGNSPYSGRGNSPYSGRGNSPYSGRGNSPYSGRGNSPYSGRGNSP
YSGRGNSPYSGRGNSPYSGRGNSPYSGRGNSPYSGRGNSPYSGRGNSPYSGRGNSPYSGRGNSPY
SGRGNSPYSGRGNSPYSGRGNSPYSGRGNSPYSGT-

**GRGNSPWS**

MRGSHHHHHHGSRSENLYFQGRSEFDPGRGNSPWSGRGNSPWSGRGNSPWSGRGNSPWSGRG
NSPWSGRGNSPWSGRGNSPWSGRGNSPWSGRGNSPWSGRGNSPWSGRGNSPWSGRGNSPWSG
RGNSPWSGRGNSPWSGRGNSPWSGRGNSPWSGRGNSPWSGRGNSPWSGRGNSPWSGRGNSPW
SGRGNSPWSGRGNSPWSGRGNSPWSGRGNSPWSGRGNSPWSGT-

**GRGNSPFS**

MRGSHHHHHHGSRSENLYFQGRSEFDPGRGNSPFSGRGNSPFSGRGNSPFSGRGNSPFSGRGNSP
FSGRGNSPFSGRGNSPFSGRGNSPFSGRGNSPFSGRGNSPFSGRGNSPFSGRGNSPFSGRGNSPFSG
RGNSPFSGRGNSPFSGRGNSPFSGRGNSPFSGRGNSPFSGRGNSPFSGRGNSPFSGRGNSPFSGRG
NSPFSGRGNSPFSGRGNSPFSGRGNSPFSGT-

**GRGNSPAS**

MRGSHHHHHHGSRSENLYFQGRSEFDPGRGNSPASGRGNSPASGRGNSPASGRGNSPASGRGNS
PASGRGNSPASGRGNSPASGRGNSPASGRGNSPASGRGNSPASGRGNSPASGRGNSPASGRGNSP
ASGRGNSPASGRGNSPASGRGNSPASGRGNSPASGRGNSPASGRGNSPASGRGNSPASGRGNSPA
SGRGNSPASGRGNSPASGRGNSPASGRGNSPASGT-

**GKGNSPYS**

MRGSHHHHHHGSRSENLYFQGRSEFDPGKGNSPYSGKGNSPYSGKGNSPYSGKGNSPYSGKGN
SPYSGKGNSPYSGKGNSPYSGKGNSPYSGKGNSPYSGKGNSPYSGKGNSPYSGKGNSPYSGKGN
SPYSGKGNSPYSGKGNSPYSGKGNSPYSGKGNSPYSGKGNSPYSGKGNSPYSGKGNSPYSGKGN
SPYSGKGNSPYSGKGNSPYSGKGNSPYSGKGNSPYSGT-

**GRGASPYA**

MRGSHHHHHHGSRSENLYFQGRSEFDPGRGASPYAGRGASPYAGRGASPYAGRGASPYAGRGA
SPYAGRGASPYAGRGASPYAGRGASPYAGRGASPYAGRGASPYAGRGASPYAGRGASPYAGRG
ASPYAGRGASPYAGRGASPYAGRGASPYAGRGASPYAGRGASPYAGRGASPYAGRGASPYAGR
GASPYAGRGASPYAGRGASPYAGRGASPYAGRGASPYAGT-

**Supplementary Figures:**

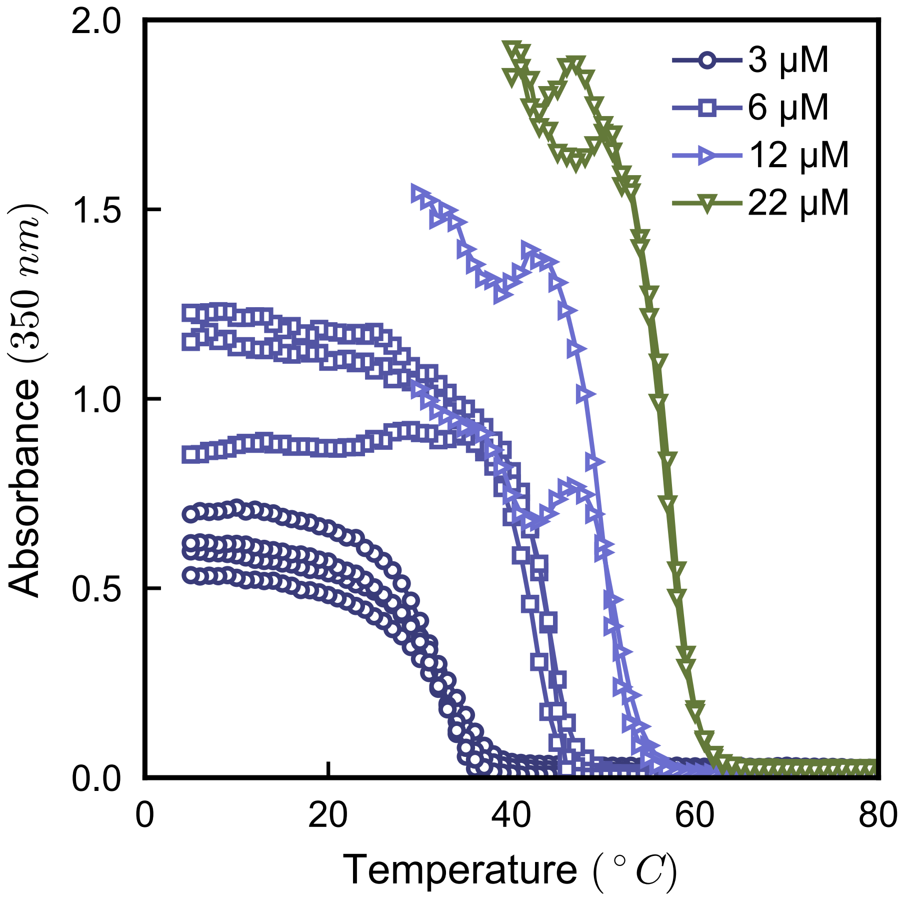


**Fig S1.** Turbidity measurements for the WT at 4 different concentrations in PBS. The transition temperatures measured are then converted into the partial phase diagram shown in Fig 1.

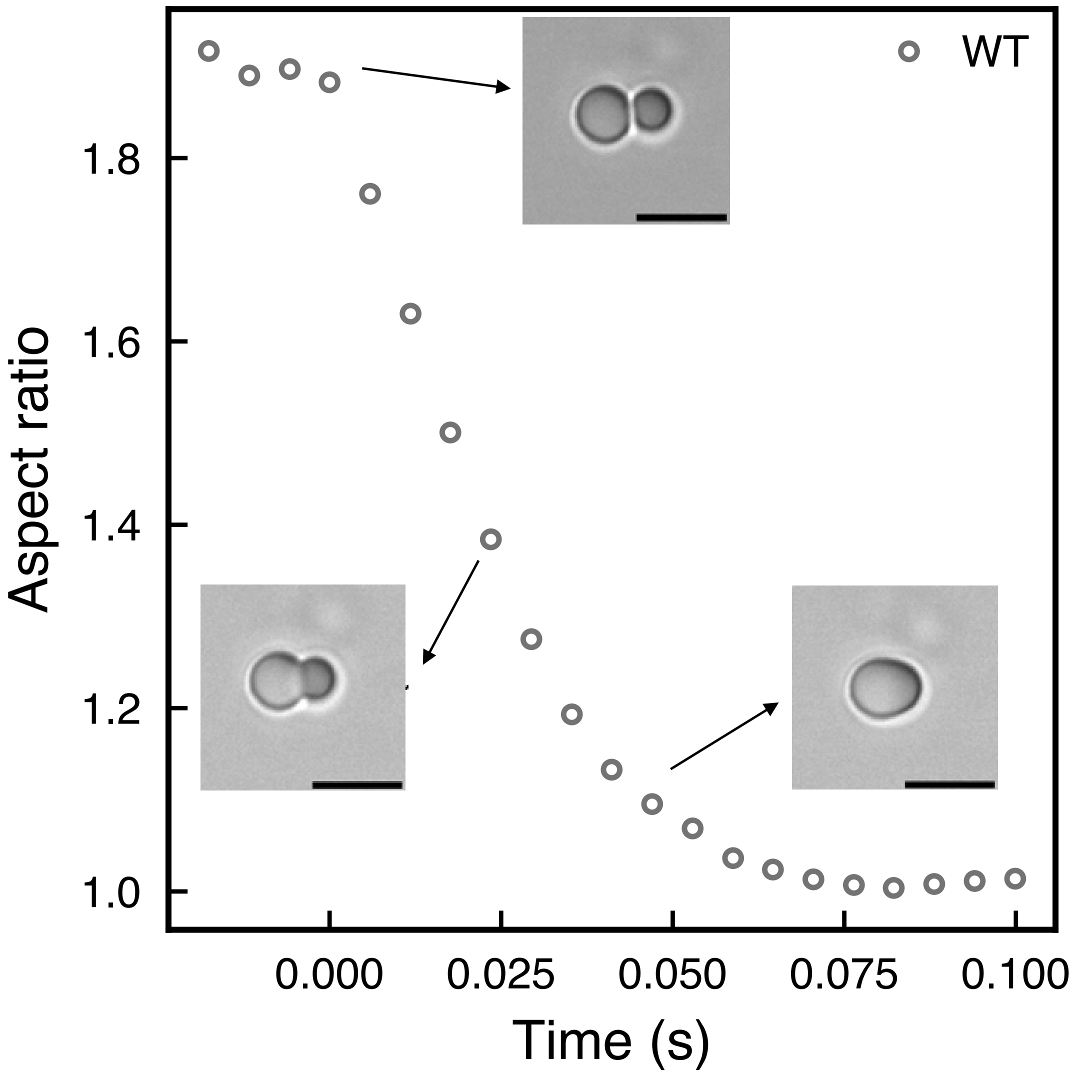


**Fig S2.** Plot of aspect ratio of fusing droplets relaxing exponentially to a sphere. The droplets are of the WT sequence. Scale bar is 5 μm.

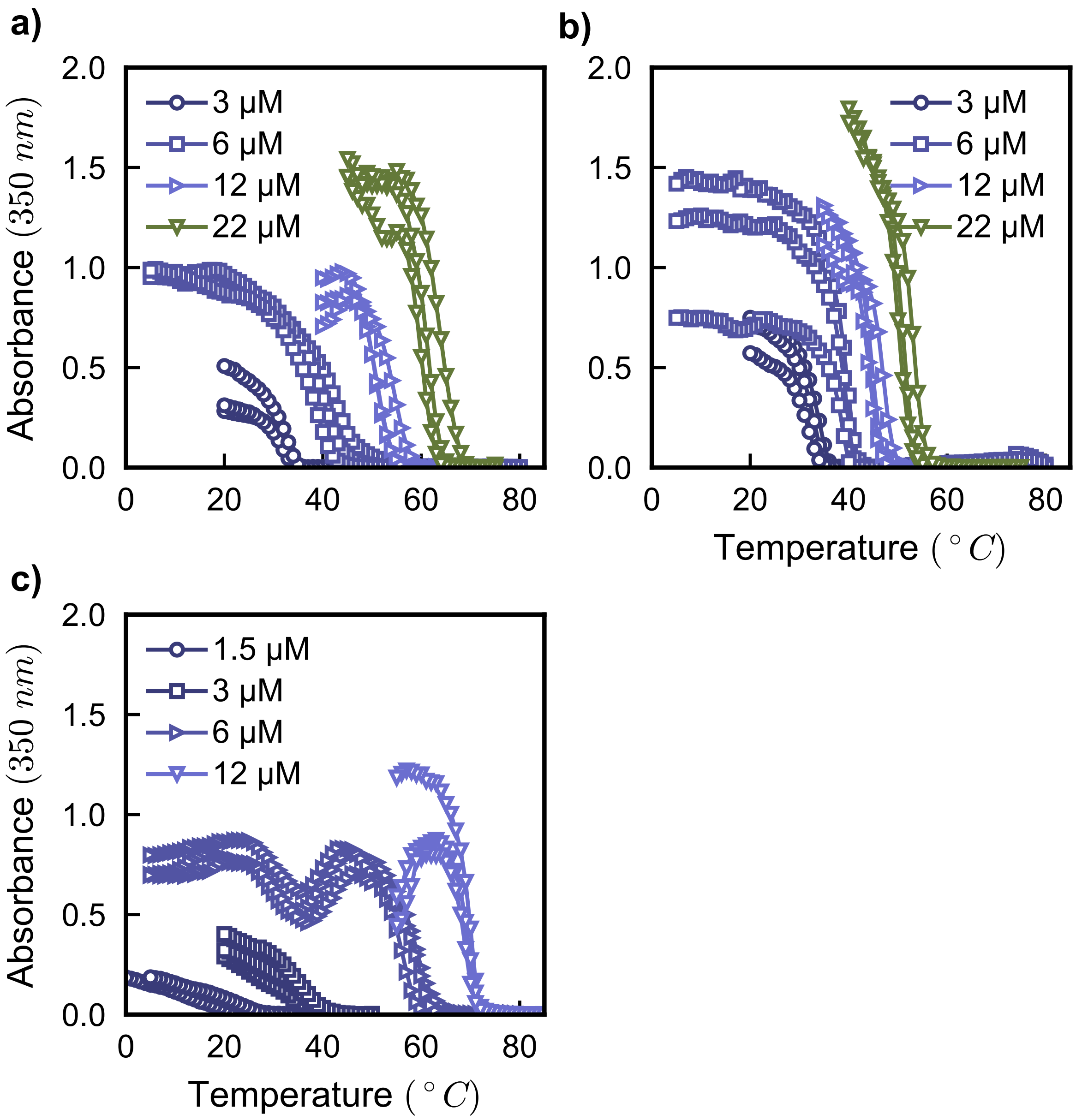


**Fig S3.** Turbidity experiments for (a) <u>A</u>R<u>A</u>DSPYS, (b) <u>S</u>R<u>S</u>DSPYS, and (c) GRGD<u>V</u>PYS conducted at 4 different concentrations in PBS.

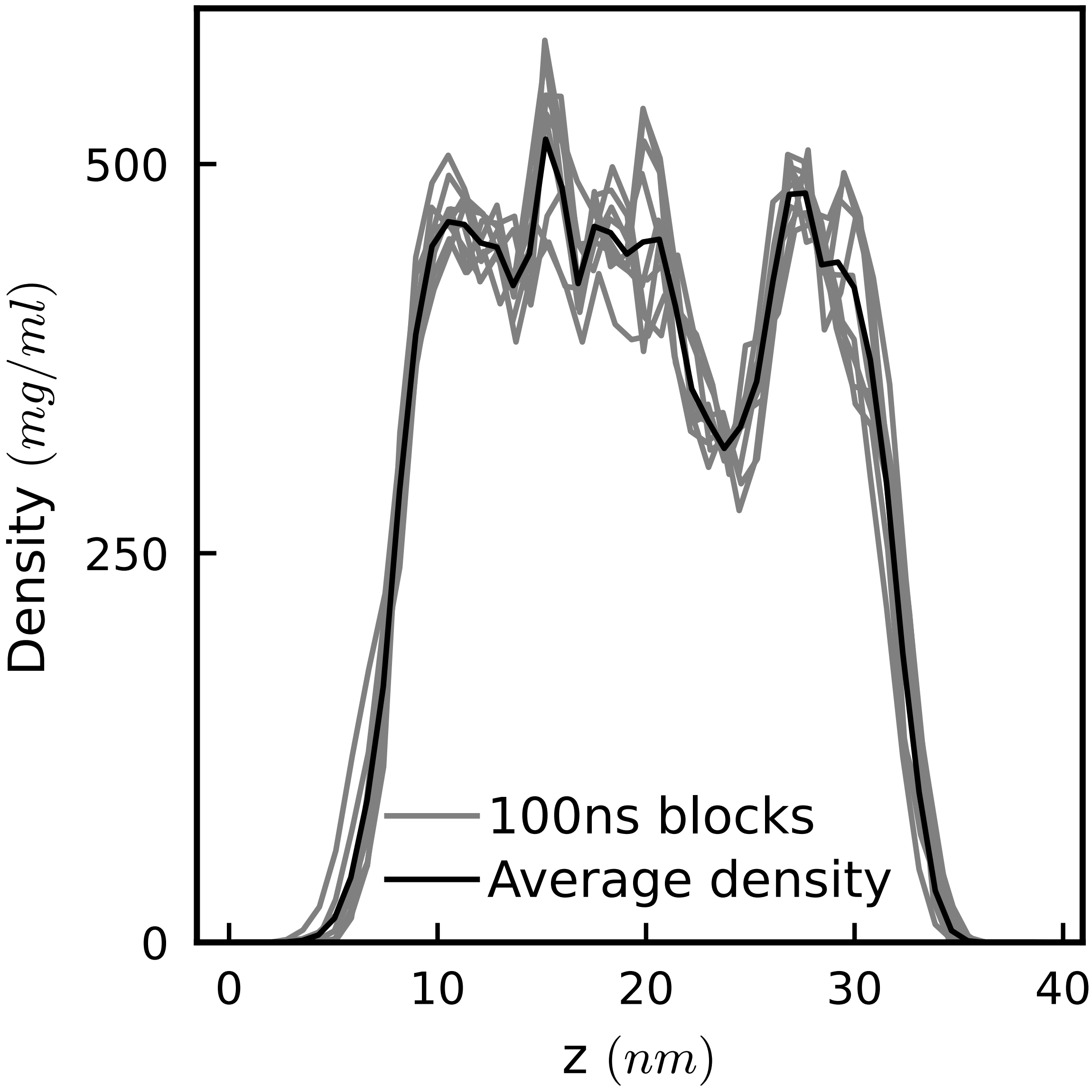


**Fig S4.** Density profile of the WT A-IDP calculated from atomistic slab simulations. The lines in gray represent calculated densities split into 10 blocks of 100 ns each, e.g., 0-100 ns, 100-200 ns and so on, while the black line represents the averaged density profile over the entire 1 µs trajectory highlighting that the protein density remains stable through the course of the simulation.

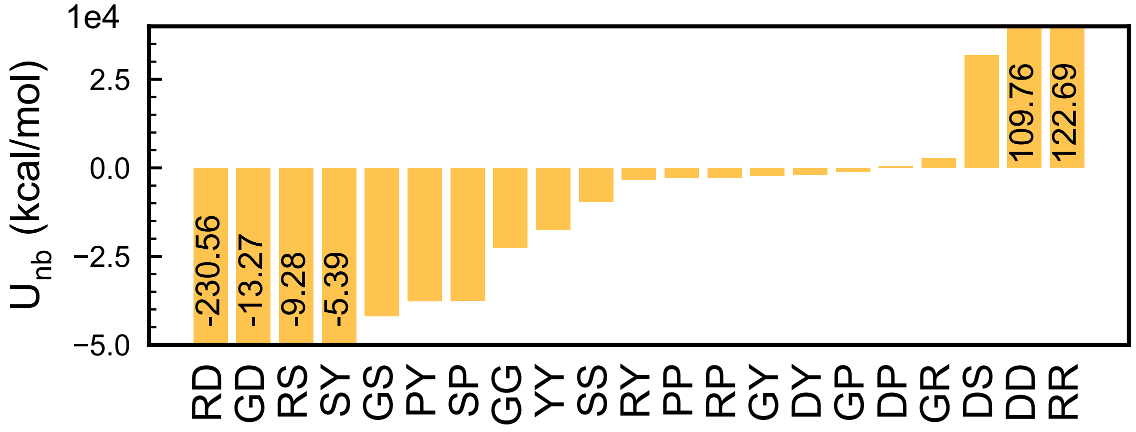


**Fig S5.** Total pairwise non-bonded potential energy (van der Waals + electrostatic) between residue pairs present in the WT A-IDP estimated from atomistic simulation trajectories. Negative values indicate more favorable interactions between the residue pairs, while positive values indicate repulsion between residue pairs. The numerical values are shown inside the bars for residue pairs with the higher energies.

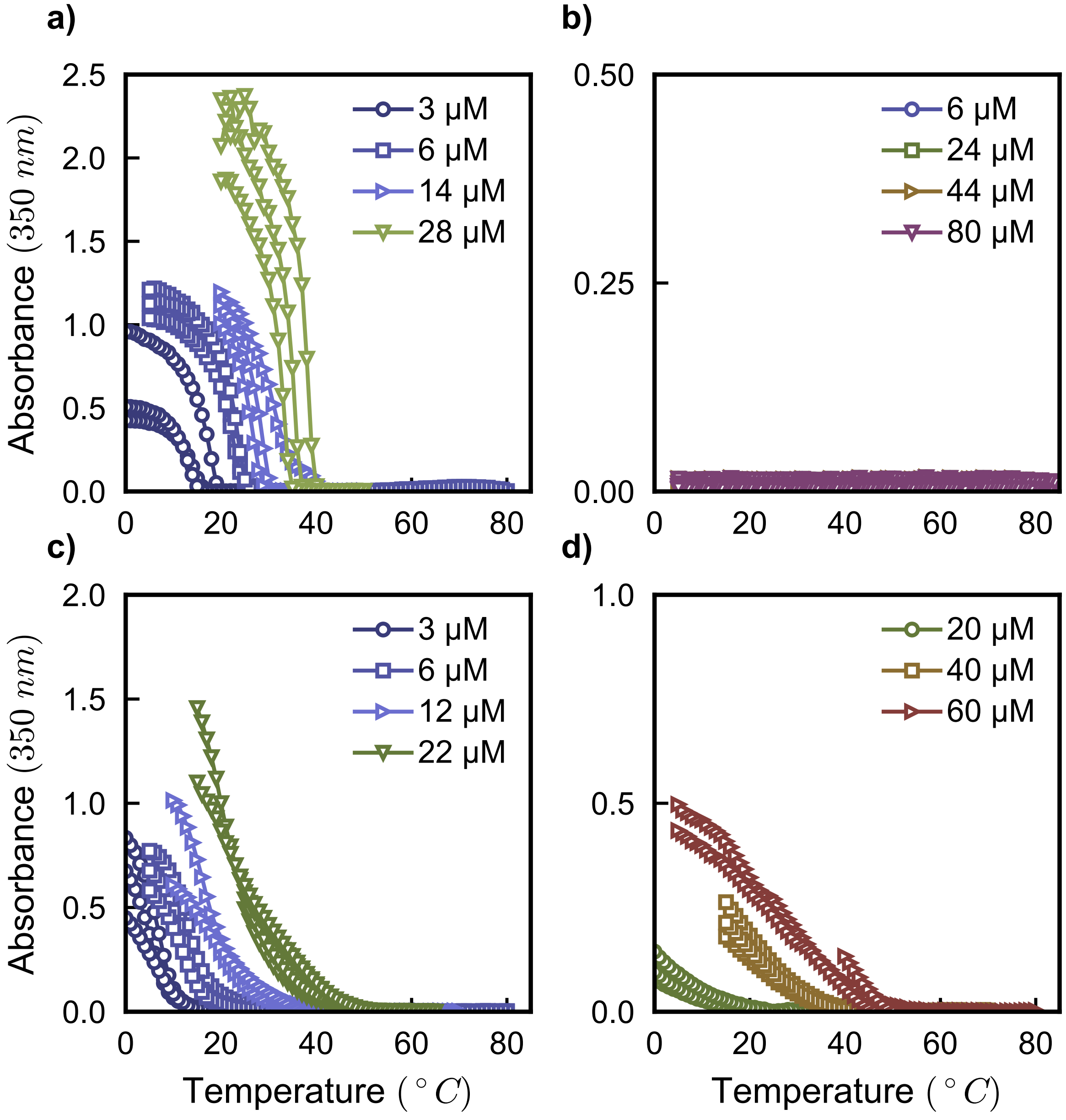


**Fig S6.** Turbidity experiments for (a) GQGNSPYS, (b) GQGDSPYS, (c) GRGNPSYS, and (d) GKGNSPYS at different concentrations in PBS. Since the GKGNSPYS variant only phase separates at higher concentrations, the measurements are shown from 20 μM onwards. The GQGDSPYS variant shows no measurable transition even up to 80 μM.

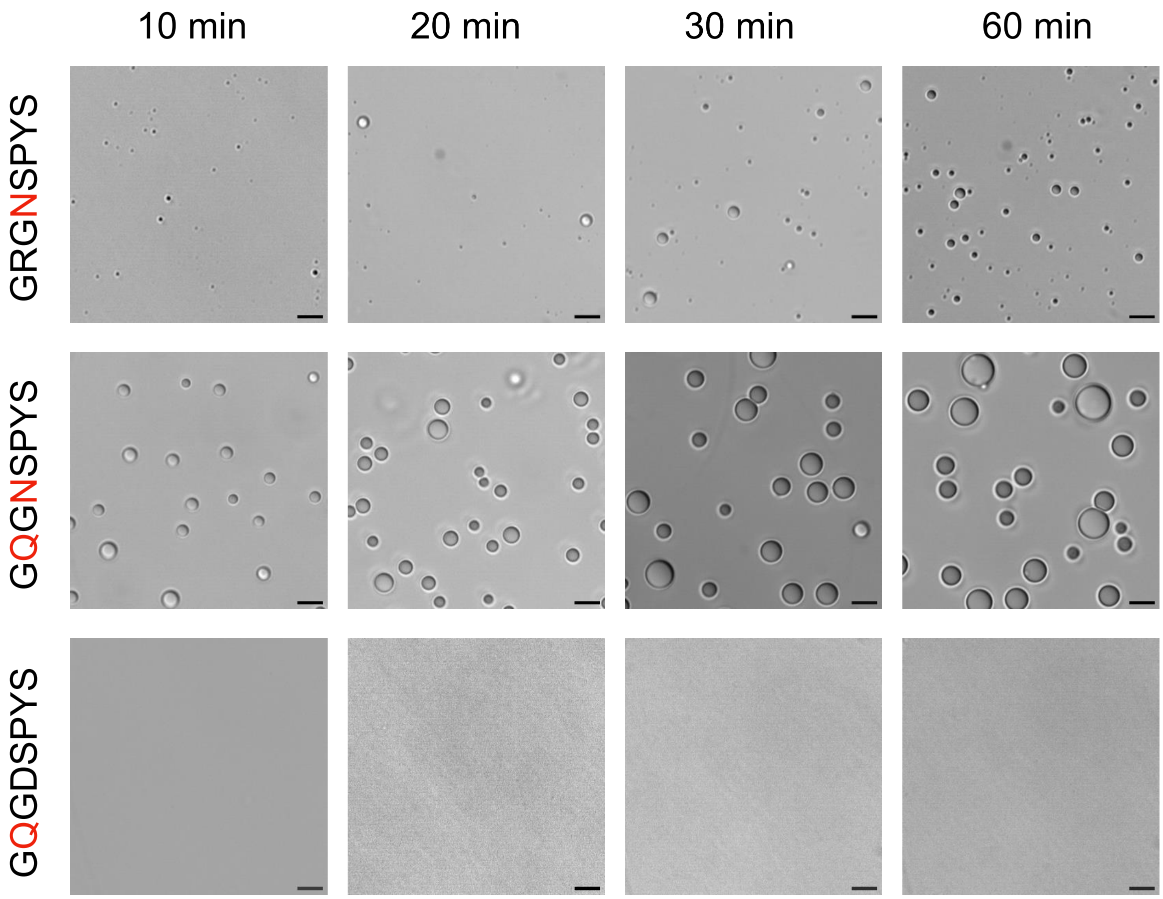


**Fig S7.** Microscopy images showing comparison of droplet growth over time for cationic sequence (GRG<u>N</u>SPYS), neutral sequence (G<u>Q</u>G<u>N</u>SPYS), and anionic sequence (G<u>Q</u>GDSPYS). The cationic sequence with tyrosine shows slower growth of droplets as compared to the neutral sequence. (Scale bar: 5 µm)

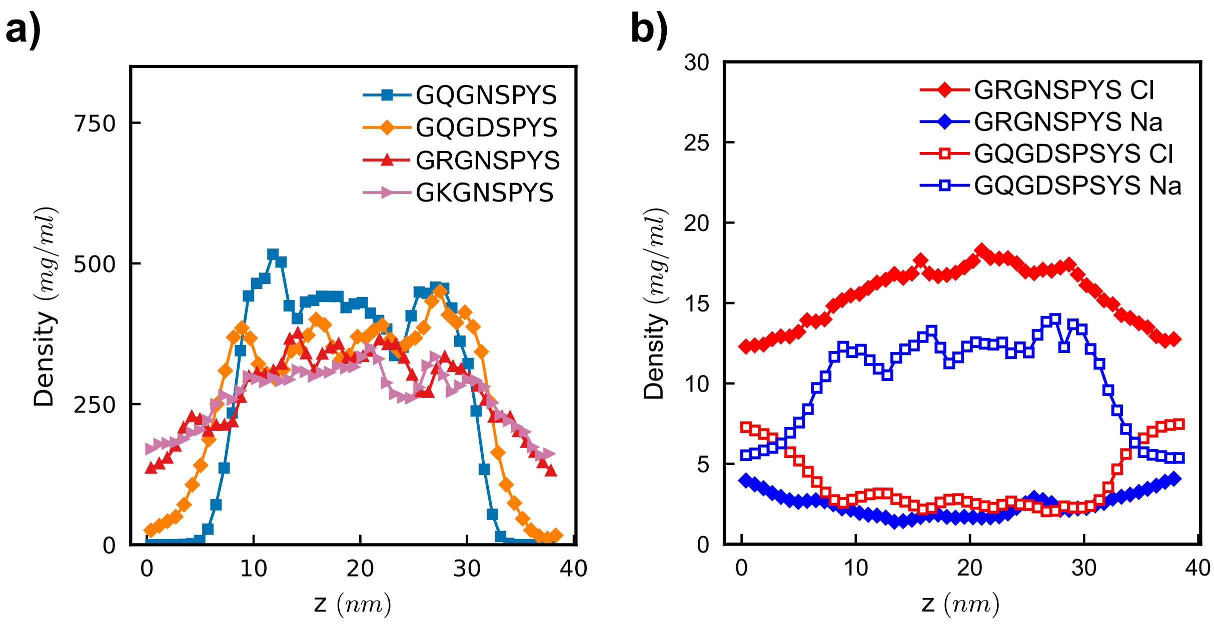


**Fig S8.** (a) Protein density profile as a function of z dimension for variants presented in Figure 3 (GQGNSPYS, GQGDSPYS, GRGNSPYS, and GKGNSPYS). Densities are averaged over the entire 1 µs atomistic slab trajectories. (b) Ion concentrations as a function of z dimension in the atomistic slab for the two variants with net charge, GQGDSPYS (empty symbols) and GRGNSPYS (filled symbols). Ion concentrations are averaged over the entire 1 µs atomistic slab trajectories.

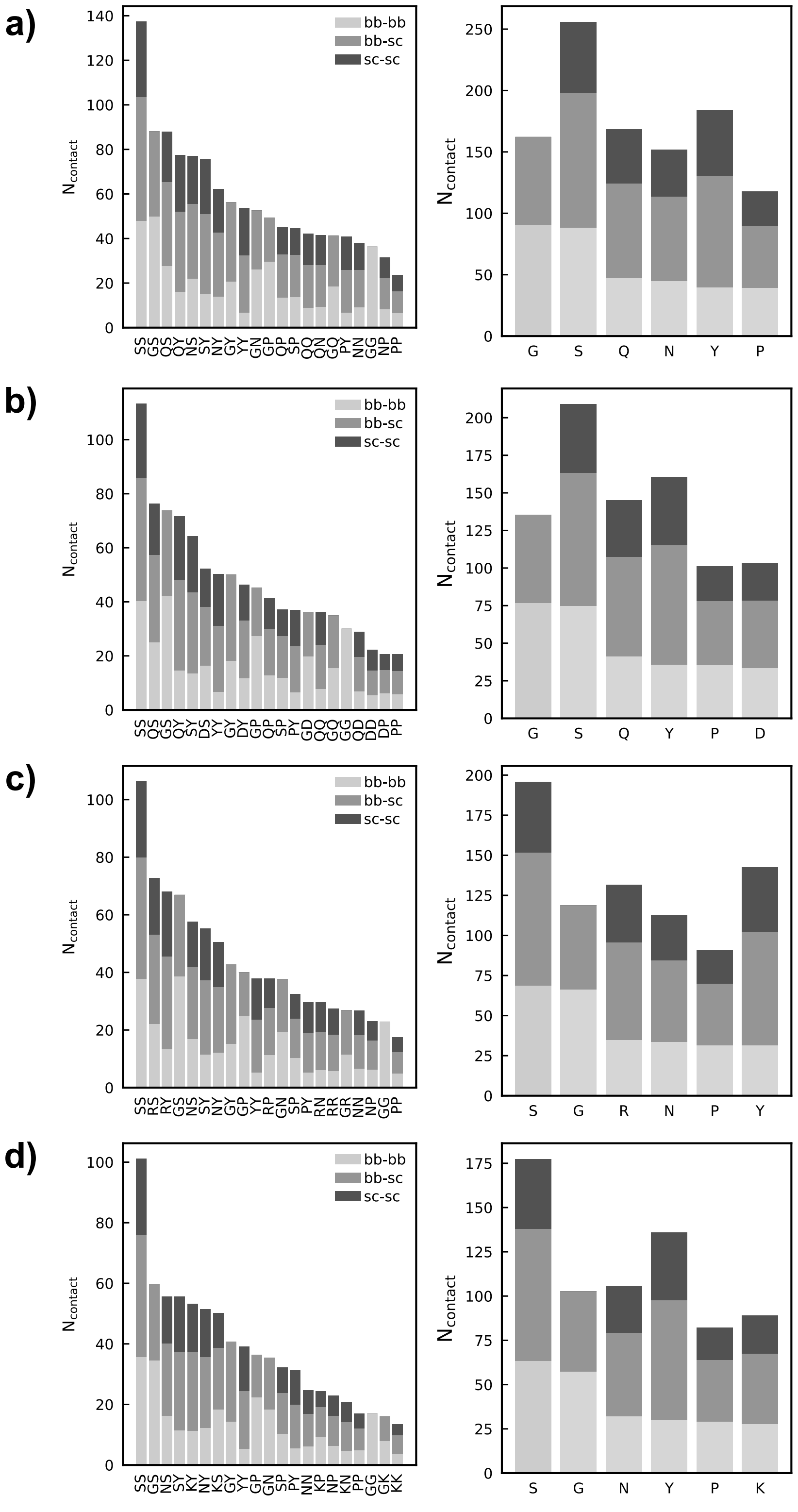


**Fig S9.** Average residue pairwise contacts (left) and per-residue contacts (right) for the (a) GQGNSPYS, (b) GQGDSPYS, (c) GRGNSPYS, and (d) GKGNSPYS variants, estimated from the atomistic slab trajectories. Both residue pair and per-residue contacts are decomposed into backbone-backbone, backbone-sidechain, and sidechain-sidechain. (Data is not normalized by residue abundance in the sequences.)

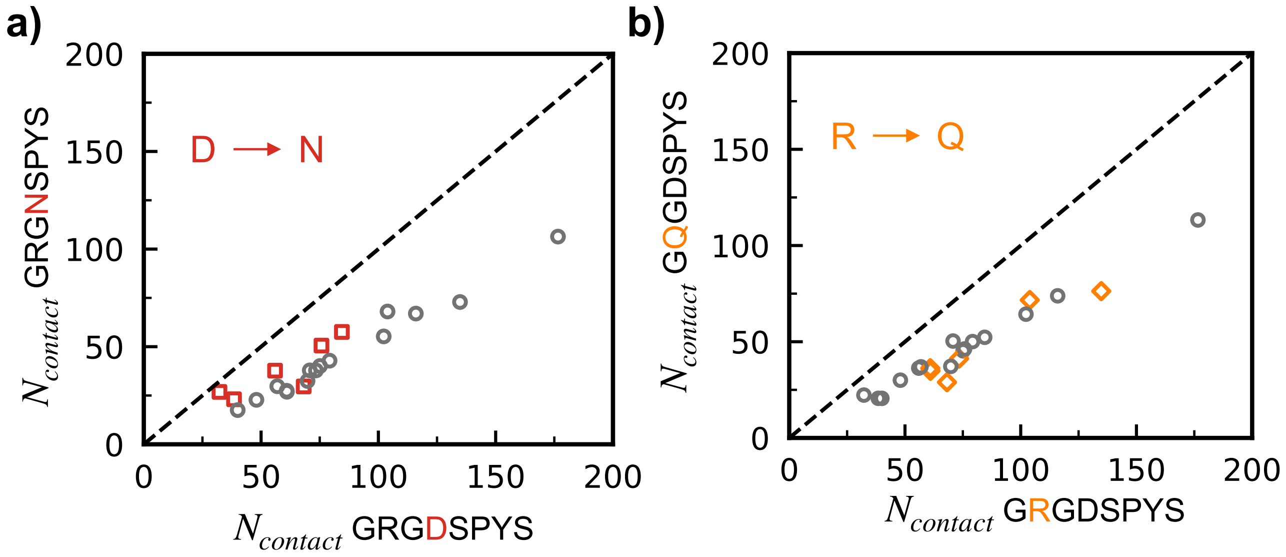


**Fig S10.** Average residue pairwise contacts for the (a) GRGNSPYS and (b) GQGDSPYS variants with respect to WT. Residue pairs not involving mutated residues are shown as dark gray circles while residue pairs involving the mutated residues are shown as red squares (D to N) and orange diamonds (R to Q), respectively.

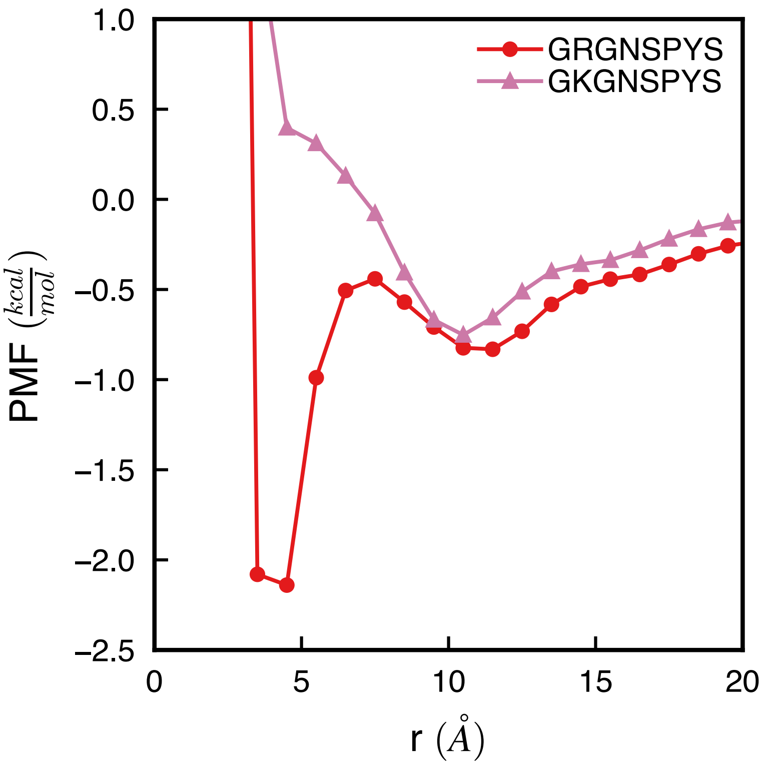


**Fig S11.** Potential of mean force between atoms of the cationic residue (arginine and lysine, respectively) and the atoms of the aromatic ring of tyrosine for the GRGNSPYS and GKGNSPYS variants. PMFs are estimated through calculation of the radial distribution function between the atom groups. Hydrogen atoms are excluded from the analysis.

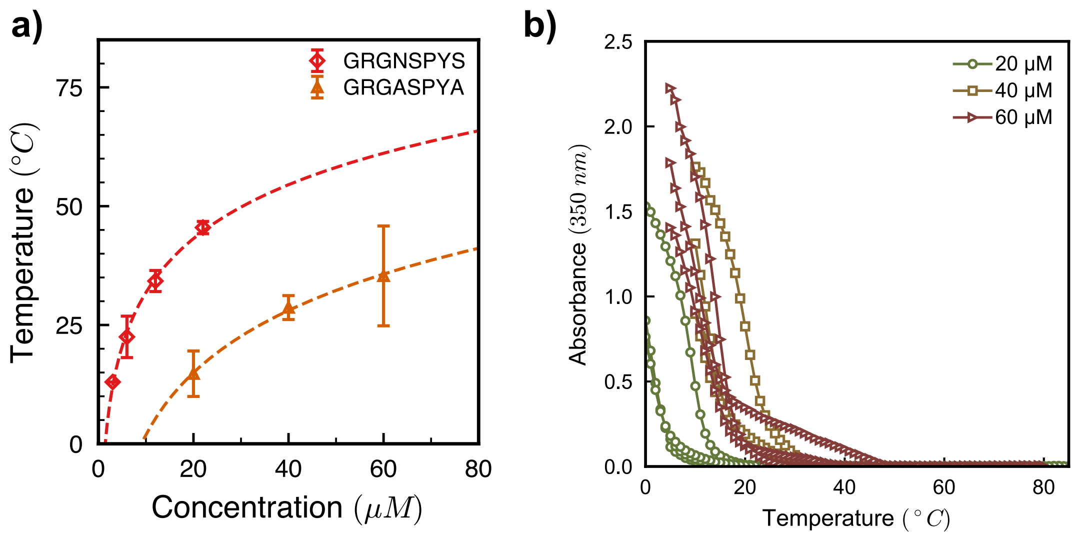


**Fig S12.** Partial phase diagram (a) and turbidity measurements (b) at different concentrations in PBS for the GRGASPYA variant.

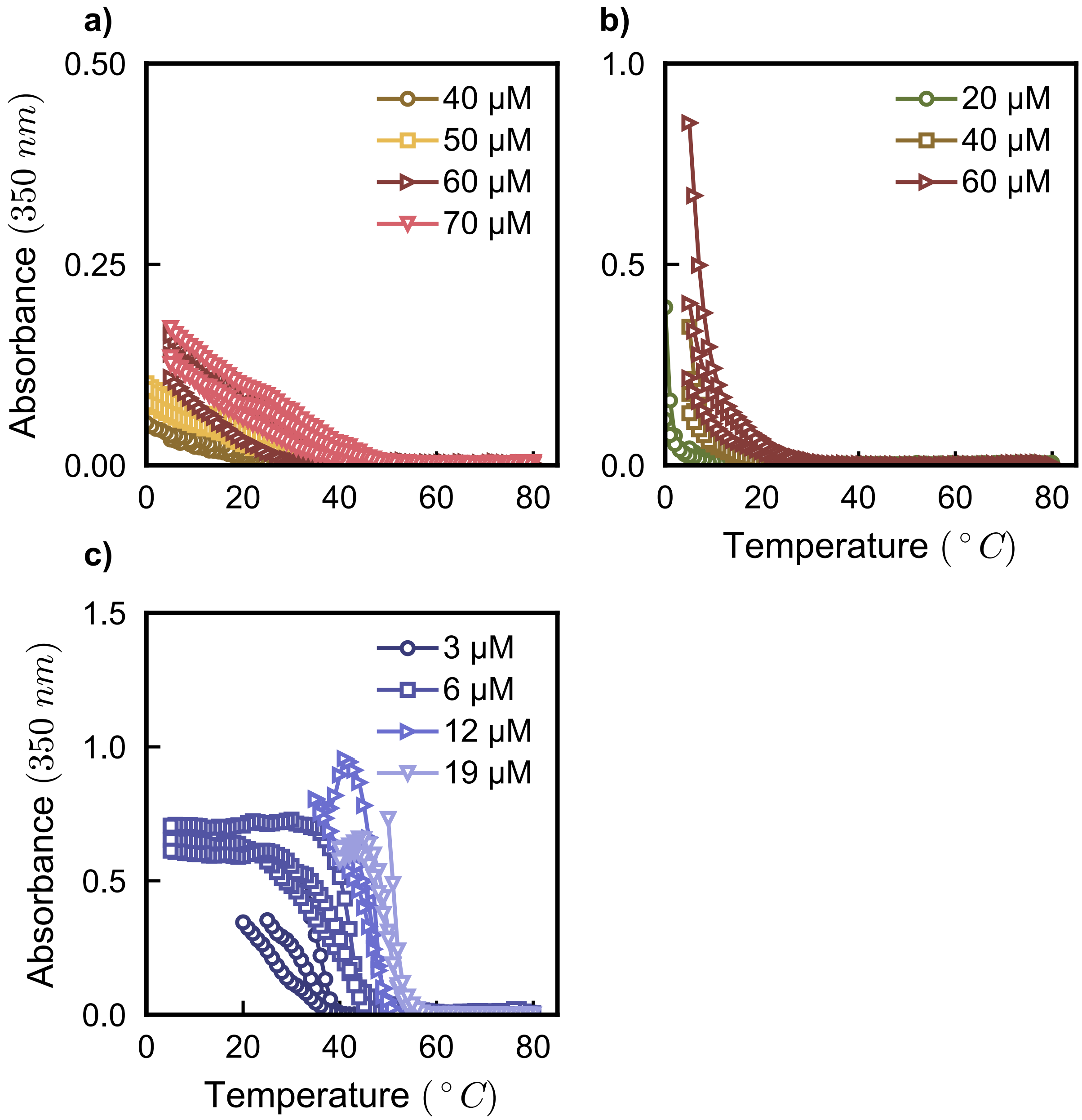


**Fig S13.** Turbidity measurements at different concentrations in PBS for the (a) GRGNSPAS, (b) GRGNSPFS, and (c) GRGNSPWS variants. GRGNSPAS and GRGNSPFS undergo LLPS at higher concentrations; therefore, for clarity, turbidity assays at lower concentrations where a transition is not observed are omitted.

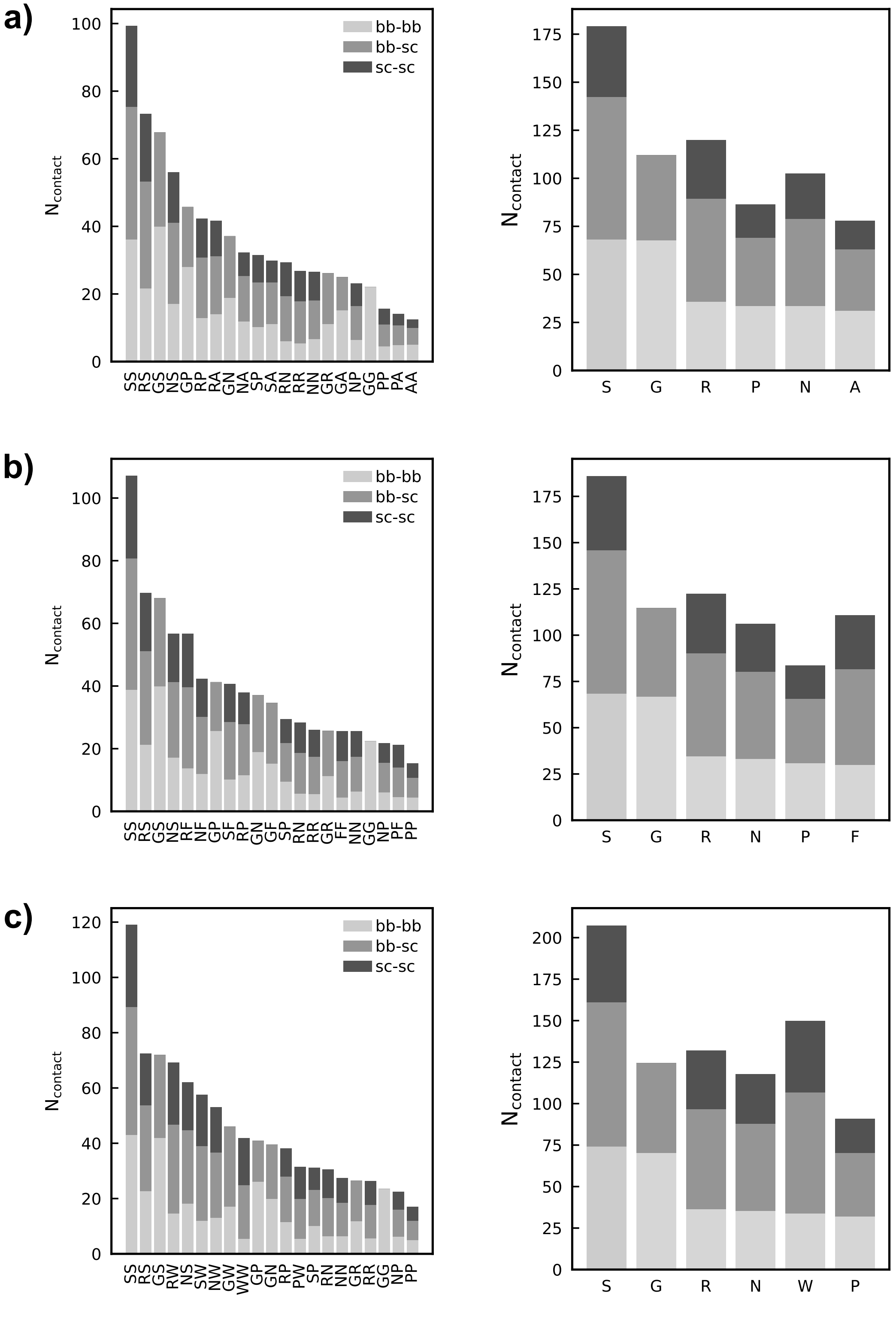


**Fig S14.** Average residue pair contacts (left) and per-residue contacts (right) for the (a) GRGNSPAS, (b) GRGNSPFS, and (c) GRGNSPWS variants estimated from the atomistic slab trajectories. Both residue pair and per-residue contacts are decomposed into backbone-backbone, backbone-sidechain, and sidechain-sidechain. (Data is not normalized by residue abundance in the sequences.)

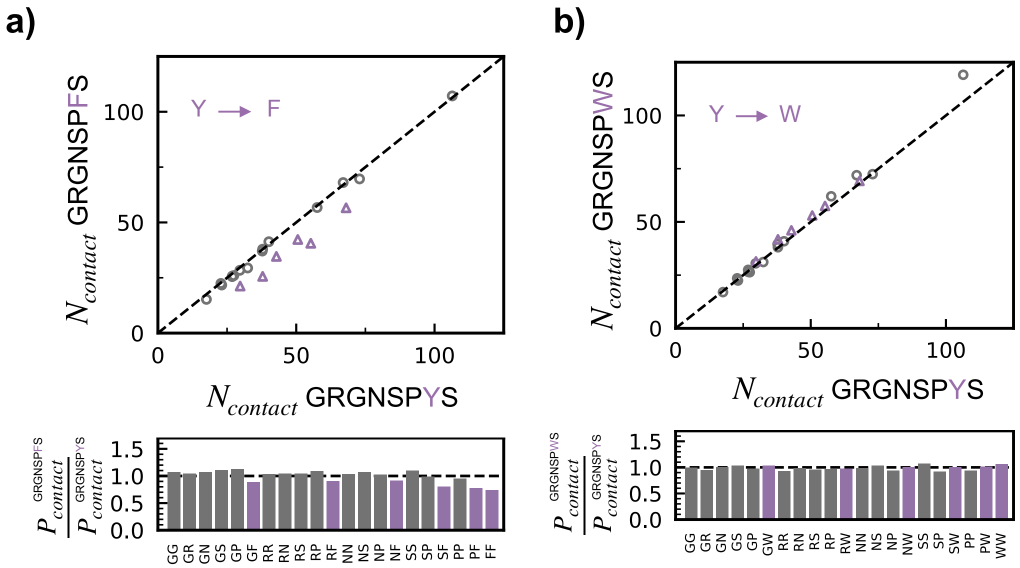


**Fig S15.** Average residue pairwise contacts for the (a) GRGNSPFS and (b) GRGNSPWS variants with respect to GRGNSPYS. Upper plots: Residue pairs not involving mutated residues are shown as dark gray circles while residue pairs involving the mutated residues are shown as purple triangles. Lower plots: Ratio between the probability of contact formation for the GRGNSPFS and GRGNSPWS variants to that of the GRGNSPYS variant.

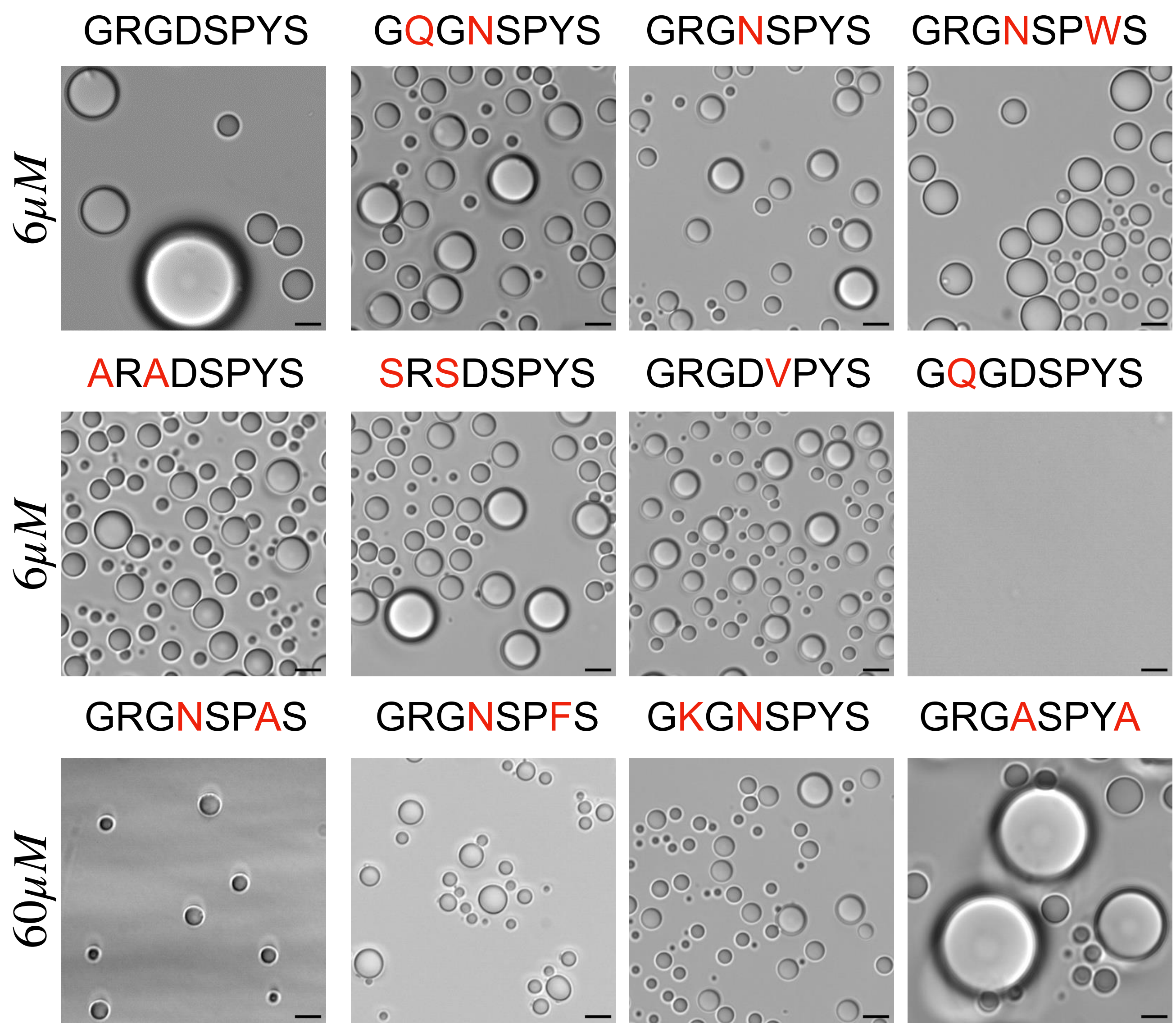


**Fig S16.** Microscopy images for different mutants after 24 hrs of phase separation, showing regular spherical droplets and no signs of fibrillization or aggregation. (Scale bar: 5 μm)

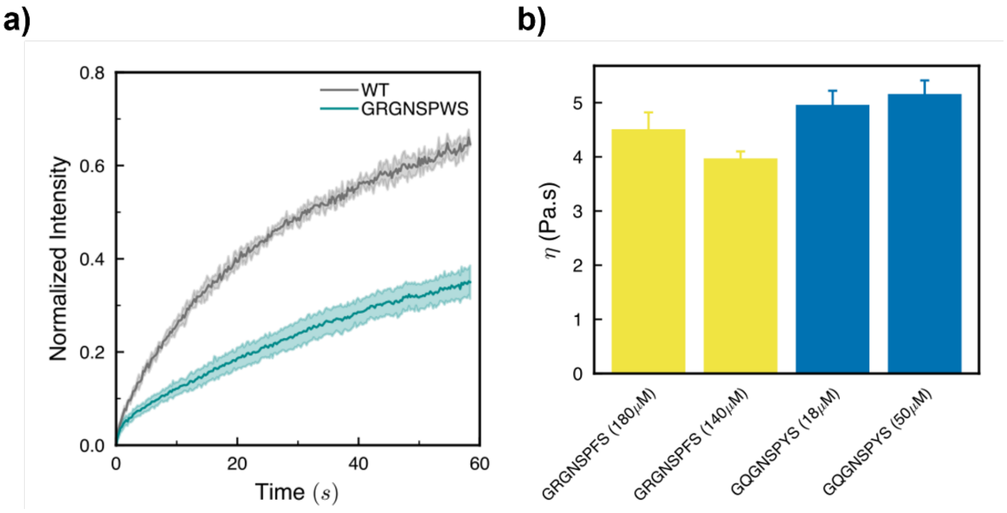


**Fig S17** (a) FRAP of WT and GRGNSPWS. FRAP measurements show a significant decrease in recovery rate for GRGNSPWS$_5$ compared to WT, which agrees with the microrheology results (Fig. 5) showing GRGNSPWS is more viscous than WT. For the FRAP measurements, RGG-GFP-RGG was used as a fluorescent tracer (see Supplementary Methods). b) Effect of total protein concentration on condensate viscosity, as measured by microrheology. Microrheology measurements were conducted for two sequences, GRGNSPFS and GQGNSPYS, at two concentrations each. The measured viscosities show minimal change with total protein concentration, suggesting that in our experiments, condensate viscosity is an intrinsic material property and does not depend on total protein concentration. This allows direct comparison of viscosities of sequences measured at different total protein concentration.

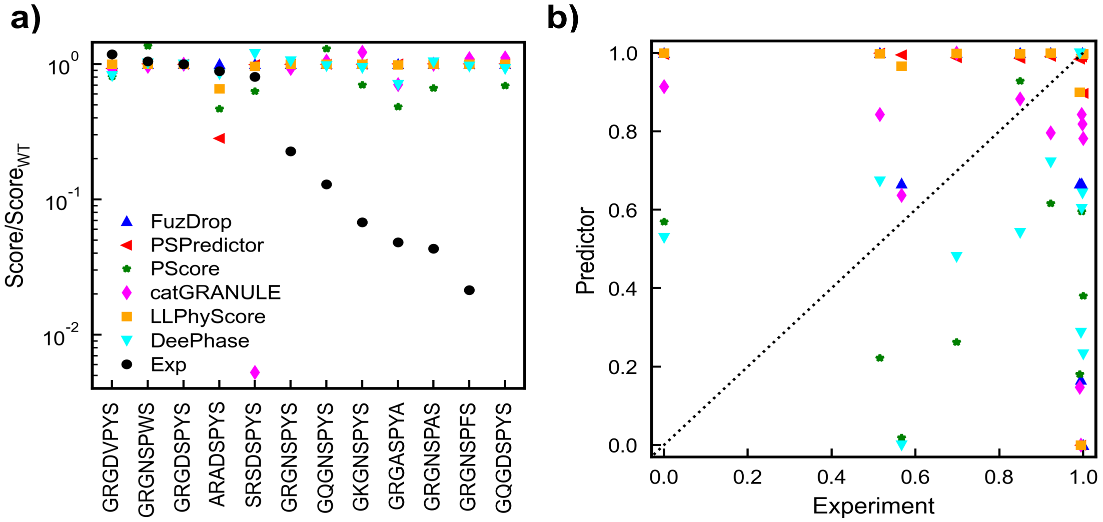


**Fig S18.** Ratio of phase separation propensity score for each sequence relative to the propensity score for WT, calculated using several online sequence-based predictors – DeePhase, PScore, PSpredictor, FuzDrop, LLPhysScore and catgranule. Experimental values are shown as black circles. All predictor values are normalized with the WT to account for different scales used by the predictors. In all cases, when the normalized score is above 1, the sequence is predicted to undergo LLPS more avidly than the WT, while values below 1 indicate a lower propensity to undergo LLPS when compared to WT. Experimental values are calculated from the saturation concentration values ($C_{sat}$) measured at 37 °C. The experimental values are represented as $C_{sat}$ of WT divided by $C_{sat}$ of variant, such that here too, a value above 1 indicates greater phase separation propensity compared to WT, whereas a value below 1 indicates lower phase separation propensity. (b) Correlation between experimental values and predictor results. Data for all data sets are normalized from 0 to 1. Symbols are the same as shown in (a) for the predictors.

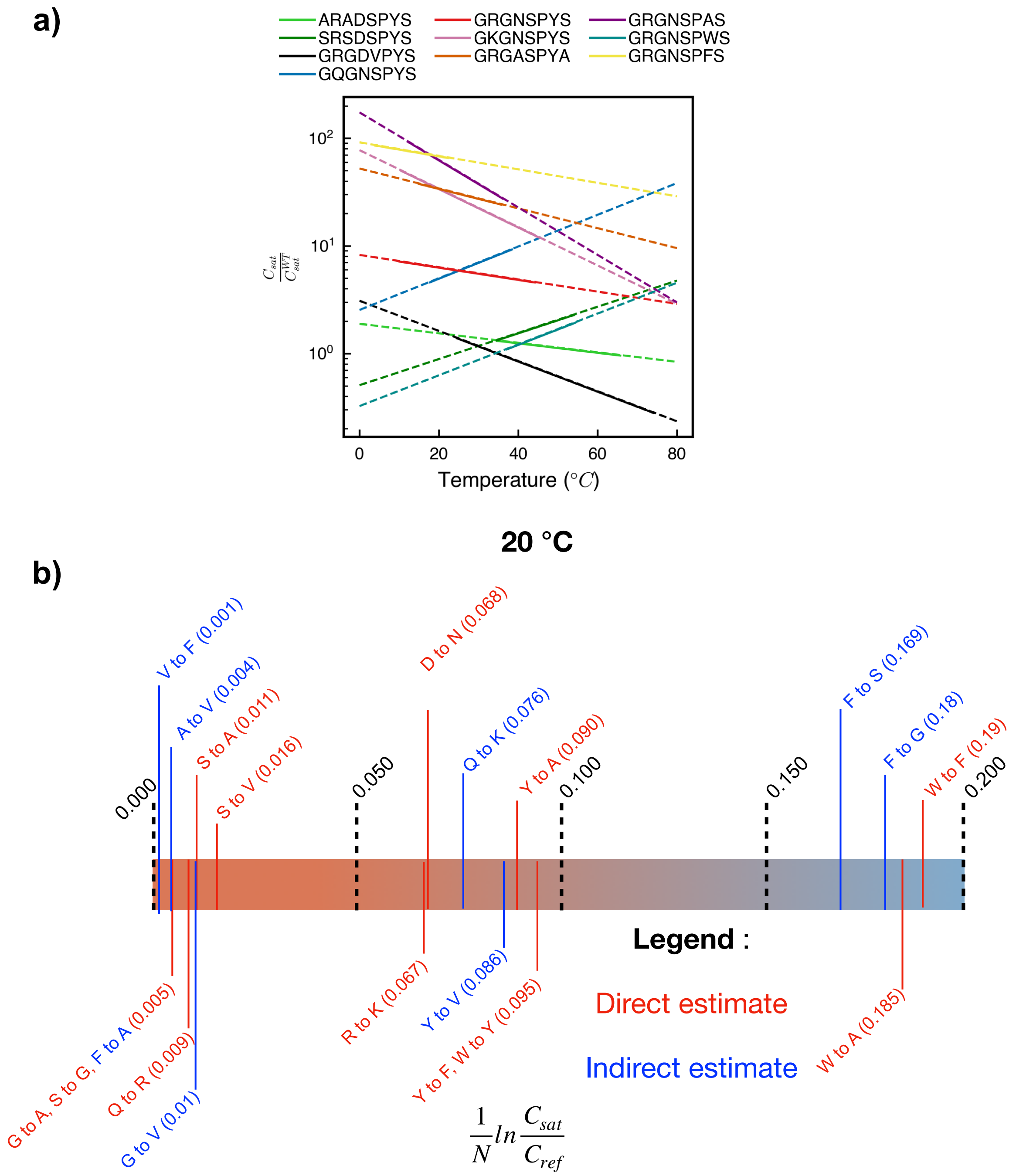


**Fig S19.** (a) Ratio of saturation concentrations ($C_{sat}$) for different sequences with respect to $C_{sat}$ of WT at different temperatures. Lines sloping down indicate that phase separation propensity with respect to the WT is enhanced at higher temperatures, whereas lines sloping upwards indicate reduction in phase separation propensity with respect to WT at higher temperatures. Solid lines indicate the temperatures at which saturation concentration was estimated using turbidimetry experiments, while dashed lines indicate the temperatures at which values were extrapolated from a logarithmic fit to the experimental binodal data. (b) Thermodynamic analysis performed for the different variants based on the estimated saturation concentrations at 20 °C. Higher values indicate reduction in phase separation propensity upon carrying out the mutation. Direct estimate refers to values that can be calculated from the experimental variants directly, whereas indirect estimate refers to values for which the mutation was not carried out in this work and the values were inferred based on data from multiple related experimental variants.

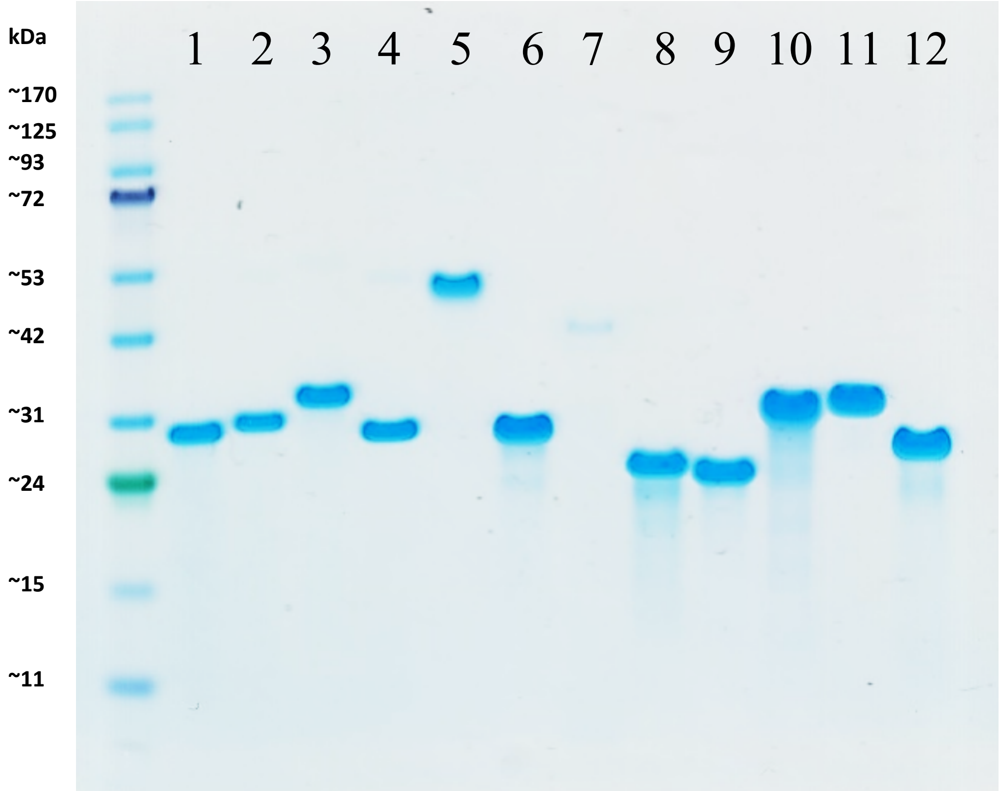


1. GRGDSPYS (WT)
2. ARADSPYS
3. SRSDSPYS
4. GRGDVPYS
5. GQGNSPYS
6. GRGNSPYS
7. GQGDSPYS
8. GRGNSPWS
9. GRGNSPFS
10. GRGNSPAS
11. GKGNSPYS
12. GRGASPYA

**Fig S20.** SDS-PAGE of all sequences showing a unique band after purification. All samples were loaded at 0.5 mg/mL and run under standard protocols. Sequences rich in Gln (Q) have been reported to form oligomers[2] which is likely the origin of the apparent doubled MW in the sequences in columns 5 and 7. The polyanionic sequence GQGDSPYS, in column 7, shows a weaker band compared with the other constructs, owing to weaker staining by the Coomassie blue stain, which interacts with positively charged amino acids. Slight variations in the MW of the constructs arise not only from variations in their theoretical MW but also likely variations in their charge/mass ratios as a result of their differential adsorption of SDS due to differences in their ionic character.

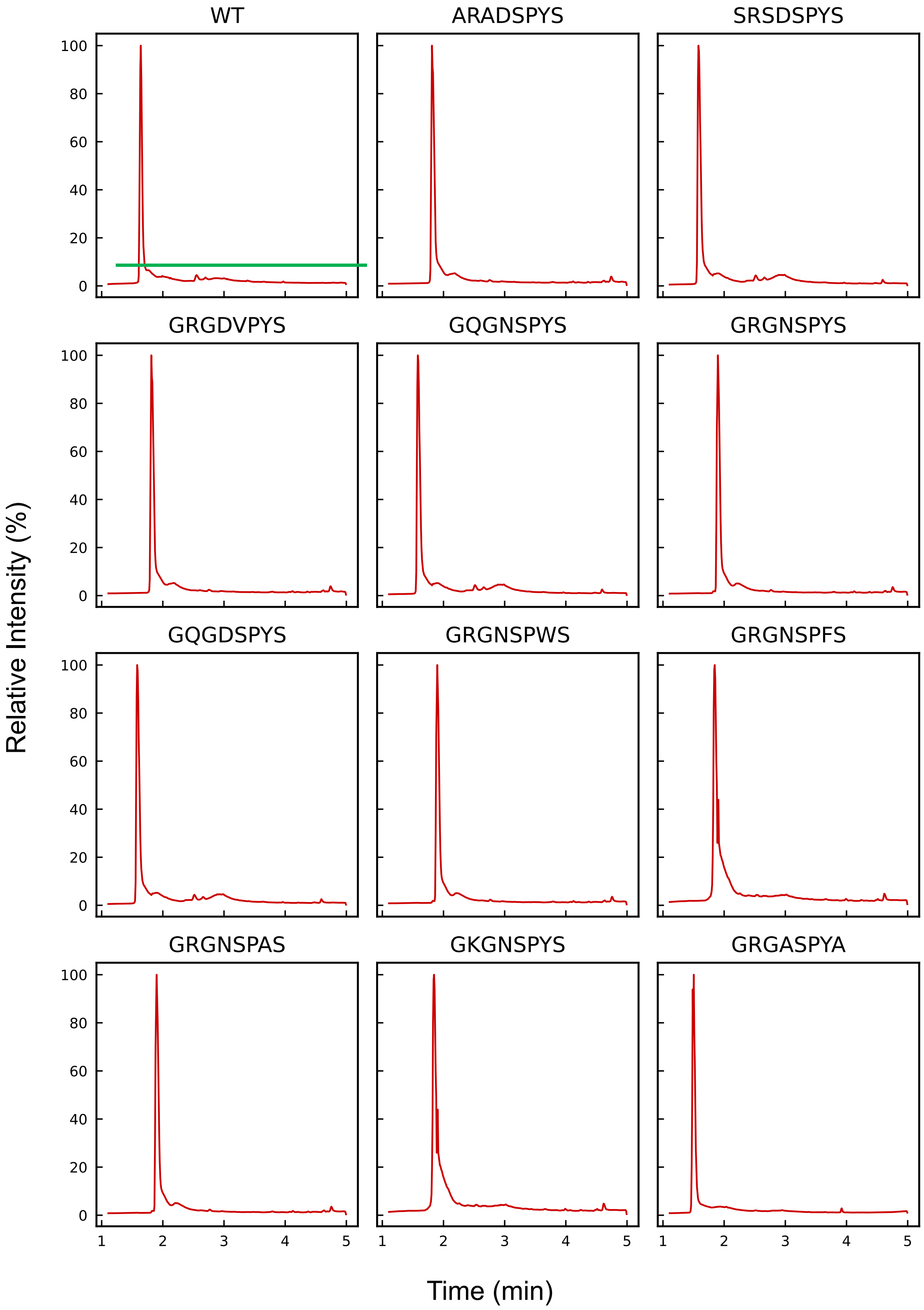


**Fig S21.** Chromatograms obtained by UPLC (XEVO). The green line in the first plot shows the area analyzed in mass spectrometry for each sequence (Fig S19).

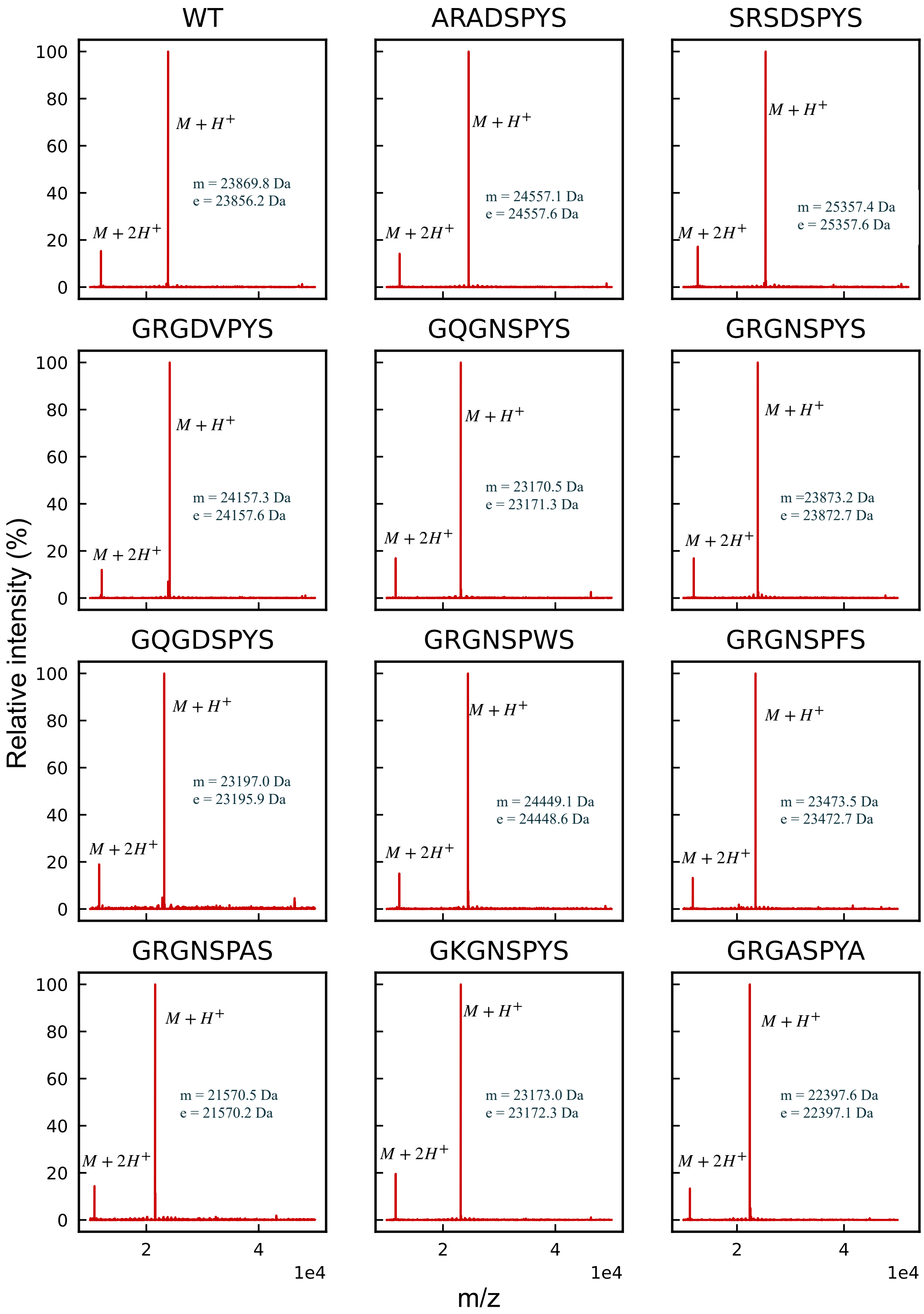


**Fig S22.** Mass spectra (XEVO) obtained from the UPLC chromatograms from Fig S18. The measured molecular weight is represented by "m" and the theoretical molecular weight is represented by "e".

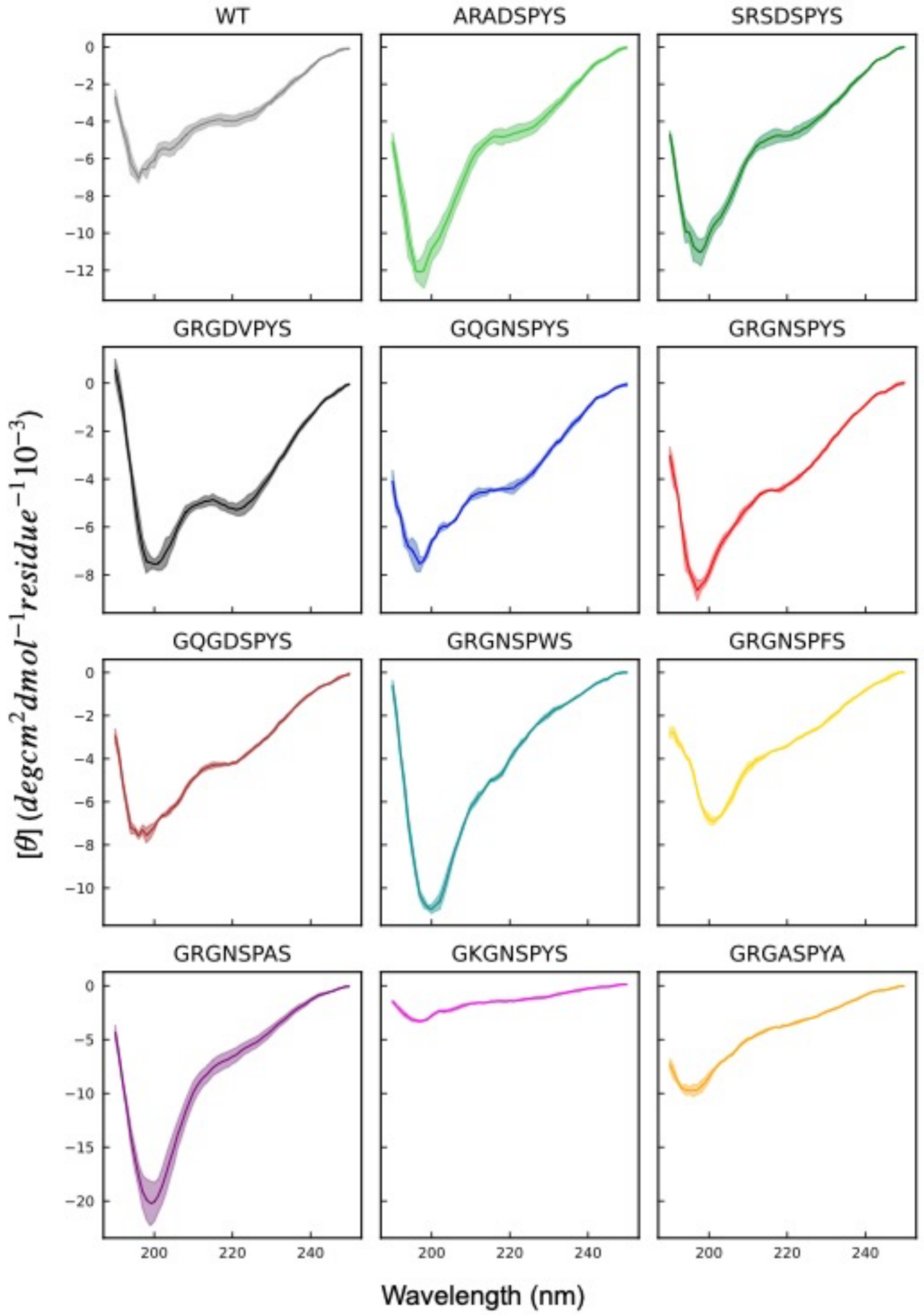


**Fig S23.** Circular dichroism (CD) spectroscopy (using a Jasco J-1500 CD spectropolarimeter, Jasco Inc., Easton, MD, USA) was conducted to characterize the secondary structure of the RLP sequences. Lyophilized samples were dissolved in DI water at pH 7.4 to a final peptide concentration of 6 µM. The CD spectra were recorded using quartz cells with a 0.2-cm optical path length. The wavelength scans were obtained at 75 °C (above the transition temperature of all the sequences at this concentration) from 190 to 250 nm and were recorded every 1 nm. The spectra show a minimum peak at ~196 nm, characteristic of random coil configurations. A detailed analysis of the structural contributions to each spectrum is delineated below in Table S4.

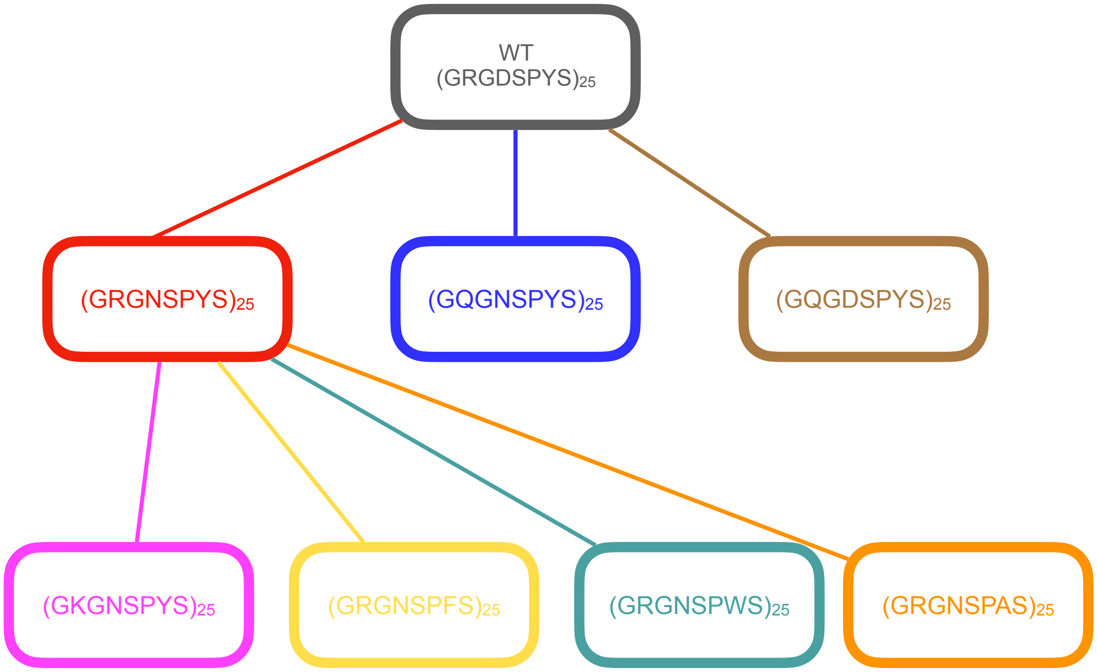


**Fig S24.** Flowchart representing the different parent and mutant sequences used in the generation of the atomistic slabs for all sequences.

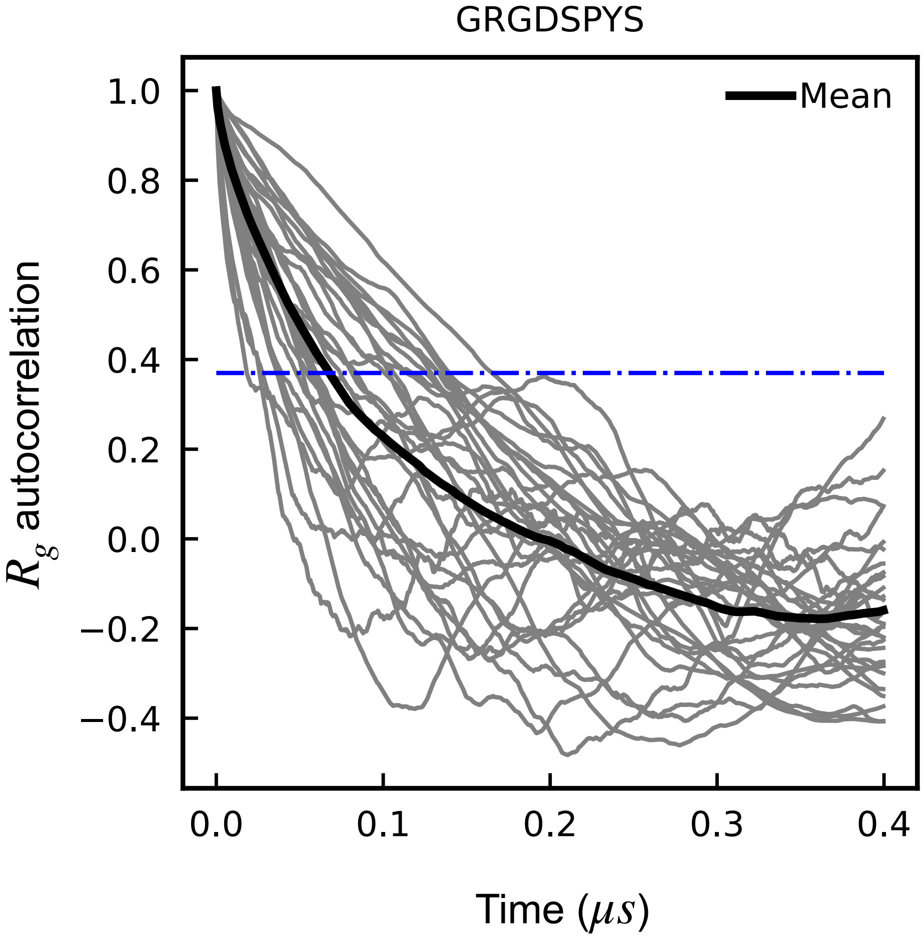


**Fig S25.** Autocorrelation of the radius of gyration for all chains in the WT atomistic slab. Individual chains are shown in grey; the mean autocorrelation value is shown in black. The horizontal dashed line in blue represents a value of 1/e.

### Supplementary Tables :

| Mutation | Author | Protein | No. of mutations | $\frac{1}{N}\ln\frac{C_{sat}}{C_{sat}^{ref}}$ |
|---|---|---|---|---|
| Y to F | Bremer et. al.[3] | hnRNPA1-LCD | 12 | 0.149 |
| | Bremer et. al.[3] | hnRNPA1-LCD | 19 | 0.157 |
| | Schuster et. al.[1] | LAF1 RGG | 10 | 0.16 |
| | Wang et. al.[4] | FUS | 27 | 0.046 |
| | Li et.al.[5] | AKAP95 | 6 | Between 0 and -0.2 |
| | This work - 20°C | A-IDP | 25 | 0.095 |
| | This work – 37°C | A-IDP | 25 | 0.094 |
| R to K | Dzuricky et. al.[6] | A-IDP | 10 | 0.20 |
| | Dzuricky et. al.[6] | A-IDP | 20 | 0.21 |
| | Bremer et. al.[3] | hnRNPA1 | 3 | 0.63 |
| | Bremer et. al.[3] | hnRNPA1 | 6 | 0.66 |
| | Brady et. al.[7] | DDX4 | 24 | >0.22 |
| | This work - 20°C | A-IDP | 25 | 0.067 |
| | This work – 37°C | A-IDP | 25 | 0.048 |
| G to A | Wang et. al.[4] | FUS | 45 | 0.01 |
| | Conicella et. al.[8] | TDP43-CTD | 1 | -1.02 |
| | This work – 20°C | A-IDP | 25 | 0.005 |
| | This work – 37°C | A-IDP | 25 | 0.002 |

**Table S1.** Thermodynamic analysis for frequently carried out mutations based on our work and findings presented in other studies. Mutations are compared based on the changes in saturation concentration with respect to reference saturation concentration upon mutation of a particular residue.

| | GRGDSPYS | | ARADSPYS | | SRSDSPYS | | GRGDVPYS | | GQGNSPYS | | GRGNSPYS | |
|---|---|---|---|---|---|---|---|---|---|---|---|---|
| | Theor | AAA | Theor | AAA | Theor | AAA | Theor | AAA | Theor | AAA | Theor | AAA |
| **Ala** | 0.0% | 1.6% | 21.8% | 20.7% | 0.0% | 0.9% | 0.0% | 1.3% | 0.0% | 0.4% | 0.0% | 0.8% |
| **Arg** | 12.2% | 11.2% | 12.2% | 11.9% | 12.2% | 12.8% | 12.2% | 11.7% | 1.3% | 1.5% | 12.2% | 12.3% |
| **Asx** | 11.8% | 12.2% | 11.8% | 12.1% | 11.8% | 13.0% | 11.8% | 11.8% | 11.8% | 12.2% | 11.8% | 12.1% |
| **Cys** | 0.0% | 0.0% | 0.0% | 0.0% | 0.0% | 0.0% | 0.0% | 0.0% | 0.0% | 0.0% | 0.0% | 0.0% |
| **Glx** | 1.3% | 3.6% | 1.3% | 2.7% | 1.3% | 2.5% | 1.3% | 2.5% | 12.2% | 11.8% | 1.3% | 2.1% |
| **Gly** | 24.0% | 21.2% | 2.2% | 3.1% | 2.2% | 3.1% | 24.0% | 21.9% | 23.6% | 23.8% | 23.6% | 23.0% |
| **His** | 2.6% | 3.5% | 2.6% | 3.3% | 2.6% | 3.1% | 2.6% | 3.2% | 2.6% | 3.0% | 2.6% | 3.0% |
| **Ile** | 0.0% | 1.2% | 0.0% | 0.9% | 0.0% | 0.7% | 0.0% | 0.9% | 0.0% | 0.3% | 0.0% | 0.7% |
| **Leu** | 0.4% | 2.0% | 0.4% | 1.7% | 0.4% | 1.4% | 0.4% | 1.6% | 0.4% | 0.8% | 0.4% | 1.3% |
| **Lys** | 0.0% | 0.9% | 0.0% | 0.7% | 0.0% | 0.5% | 0.0% | 0.7% | 0.0% | 0.2% | 0.0% | 0.4% |
| **Met** | 0.4% | 0.8% | 0.4% | 0.6% | 0.4% | 0.7% | 0.4% | 0.9% | 0.4% | 0.7% | 0.4% | 0.8% |
| **Phe** | 0.9% | 1.3% | 0.9% | 1.3% | 0.9% | 1.1% | 0.9% | 1.2% | 0.9% | 0.9% | 0.9% | 1.0% |
| **Pro** | 10.9% | 9.0% | 10.9% | 9.8% | 10.9% | 10.7% | 10.9% | 9.5% | 11.4% | 11.3% | 11.4% | 10.6% |
| **Ser** | 23.6% | 18.6% | 23.6% | 18.4% | 45.4% | 36.3% | 12.7% | 10.4% | 23.6% | 20.5% | 23.6% | 19.3% |
| **Thr** | 0.4% | 1.3% | 0.4% | 1.1% | 0.4% | 0.8% | 0.4% | 1.1% | 0.4% | 0.6% | 0.4% | 0.8% |
| **Trp** | 0.0% | 0.0% | 0.0% | 0.0% | 0.0% | 0.0% | 0.0% | 0.0% | 0.0% | 0.0% | 0.0% | 0.0% |
| **Tyr** | 11.4% | 10.0% | 11.4% | 10.8% | 11.4% | 11.5% | 11.4% | 10.5% | 11.4% | 11.7% | 11.4% | 11.2% |
| **Val** | 0.0% | 1.7% | 0.0% | 1.0% | 0.0% | 0.8% | 10.9% | 10.9% | 0.0% | 0.2% | 0.0% | 0.6% |

**Table S2.** Amino acid composition of the first set of sequences. The blue shading denotes the specific mutation for each sequence. For each sequence, a comparison between the theoretical composition (Theor) and the experimentally determined composition by amino acid analysis (AAA) is shown. Acceptable values range in ±5% of error. The larger variations shown in the table originate mainly from partial amino acid destruction during the hydrolysis procedure (e.g., Ser) and also from small amounts of impurities and quantification error in the limits of the chromatography methodology[9].

| | GQGDSPYS | | GRGNSPWS | | GRGNSPFS | | GRGNSPAS | | GKGNSPYS | | GRGASPYA | |
|---|---|---|---|---|---|---|---|---|---|---|---|---|
| | Theor | AAA | Theor | AAA | Theor | AAA | Theor | AAA | Theor | AAA | Theor | AAA |
| **Ala** | 0.0% | 0.9% | 0.0% | 0.6% | 0.0% | 0.7% | 10.9% | 10.9% | 0.0% | 0.5% | 21.8% | 21.7% |
| **Arg** | 1.3% | 1.7% | 12.2% | 13.0% | 12.2% | 12.4% | 12.2% | 12.0% | 1.3% | 1.6% | 12.2% | 12.5% |
| **Asx** | 11.8% | 12.3% | 11.8% | 12.5% | 11.8% | 12.3% | 11.8% | 12.2% | 11.8% | 12.3% | 0.9% | 1.4% |
| **Cys** | 0.0% | 0.0% | 0.0% | 0.0% | 0.0% | 0.0% | 0.0% | 0.0% | 0.0% | 0.0% | 0.0% | 0.0% |
| **Glx** | 12.2% | 12.4% | 1.3% | 1.8% | 1.3% | 2.0% | 1.3% | 2.8% | 1.3% | 1.8% | 1.3% | 1.8% |
| **Gly** | 23.6% | 23.1% | 23.6% | 24.0% | 23.6% | 23.4% | 23.6% | 22.2% | 23.6% | 24.1% | 23.6% | 23.5% |
| **His** | 2.6% | 2.6% | 2.6% | 3.4% | 2.6% | 3.0% | 2.6% | 3.8% | 2.6% | 3.0% | 2.6% | 3.7% |
| **Ile** | 0.0% | 0.6% | 0.0% | 0.3% | 0.0% | 0.5% | 0.0% | 0.9% | 0.0% | 0.5% | 0.0% | 0.3% |
| **Leu** | 0.4% | 1.2% | 0.4% | 0.8% | 0.4% | 1.1% | 0.4% | 1.7% | 0.4% | 1.0% | 0.4% | 0.8% |
| **Lys** | 0.0% | 0.5% | 0.0% | 0.2% | 0.0% | 0.3% | 0.0% | 0.7% | 10.9% | 11.7% | 0.0% | 0.1% |
| **Met** | 0.4% | 0.7% | 0.4% | 0.9% | 0.4% | 0.9% | 0.4% | 0.8% | 0.4% | 0.9% | 0.4% | 0.8% |
| **Phe** | 0.9% | 1.0% | 0.9% | 0.9% | 11.8% | 12.0% | 0.9% | 1.2% | 0.9% | 1.0% | 0.9% | 0.9% |
| **Pro** | 11.4% | 10.8% | 11.4% | 11.0% | 11.4% | 10.8% | 11.4% | 10.2% | 11.4% | 10.6% | 11.4% | 10.8% |
| **Ser** | 23.6% | 19.7% | 23.6% | 19.8% | 23.6% | 19.0% | 23.6% | 17.8% | 23.6% | 18.9% | 12.7% | 10.4% |
| **Thr** | 0.4% | 0.8% | 0.4% | 0.5% | 0.4% | 0.6% | 0.4% | 0.9% | 0.4% | 0.6% | 0.4% | 0.5% |
| **Trp** | 0.0% | 0.0% | 10.9% | 9.6% | 0.0% | 0.0% | 0.0% | 0.0% | 0.0% | 0.0% | 0.0% | 0.0% |
| **Tyr** | 11.4% | 11.3% | 0.4% | 0.5% | 0.4% | 0.6% | 0.4% | 0.8% | 11.4% | 11.2% | 11.4% | 10.4% |
| **Val** | 0.0% | 0.6% | 0.0% | 0.2% | 0.0% | 0.5% | 0.0% | 1.0% | 0.0% | 0.4% | 0.0% | 0.3% |

**Table S3.** Amino composition of the second set of sequences. The blue shading denotes the specific mutation for each sequence. For each sequence, a comparison between the theoretical composition (Theor) and the amino acid analysis (AAA) is shown. Acceptable values range in ±5% of error. The larger variations shown in the table originate mainly from partial amino acid destruction during the hydrolysis procedure (e.g., Ser) and also from small amounts of impurities and quantification error in the limits of the chromatography methodology[9].

| | Helix | β-sheet | | Turn | Disordered |
|---|---|---|---|---|---|
| | | Antiparallel | Parallel | | |
| GRGDSPYS | 3 | 29.2 | 6.8 | 15.5 | 45.4 |
| ARADSPYS | 5 | 26.8 | 3.9 | 17.4 | 46.9 |
| SRSDSPYS | 5.2 | 26.7 | 4.6 | 16.8 | 46.7 |
| GRGDVPYS | 2.6 | 28.3 | 7.6 | 14.9 | 46.5 |
| GQGNSPYS | 4 | 26.7 | 6.4 | 15.1 | 47.8 |
| GRGNSPYS | 3.4 | 30.8 | 5.8 | 15.5 | 44.5 |
| GQGDSPYS | 3.5 | 28.2 | 5.9 | 15 | 47.4 |
| GKGNSPYS | 0 | 37.4 | 0 | 14.6 | 48 |
| GRGNSPWS | 3.5 | 31.8 | 2.5 | 15.3 | 47 |
| GRGNSPFS | 3.8 | 26.7 | 4.8 | 15.5 | 49.3 |
| GRGNSPAS | 8.7 | 27.3 | 0 | 18 | 45.9 |
| GRGASPYA | 4.8 | 25.6 | 4.4 | 16.4 | 48.8 |

**Table S4.** Structural analysis performed in the single spectrum analysis program from BeStSel[10-12]. The method is freely available for academic use at https://bestsel.elte.hu/index.php. The table shows the percentage of each type of structure, confirming that the disordered conformation is the primary contribution to the CD spectra. In addition, the similarities in the contributions of the different secondary conformations indicates that the amino acid substitutions did not lead to significant differences in the intrinsically disordered nature of the variants as compared to the WT sequence, GRGDSPYS.

| Sequence | Viscosity (Pa.s) |
|---|---|
| $(GRGDSPYS)_{25}$ | 11.0 ± 0.5 |
| $(ARADSPYS)_{25}$ | 11.8 ± 0.9 |
| $(SRSDSPYS)_{25}$ | 19.5 ± 2.4 |
| $(GRGDVPYS)_{25}$ | 28.9 ± 1.9 |
| $(GQGNSPYS)_{25}$ | 5.0 ± 0.3 |
| $(GRGNSPYS)_{25}$ | 8.5 ± 1.6 |
| $(GRGNSPFS)_{25}$ | 4.2 ± 0.4 |
| $(GRGNSPWS)_{25}$ | 39.7 ± 2.3 |
| $(GKGNSPYS)_{25}$ | 4.9 ± 0.2 |
| $(GRGASPYA)_{25}$ | 2.3 ± 0.6 |

**Table S5.** Viscosities measured by passive microrheology for the various polypeptide sequences tested in this study.
